# Nucleoid size scaling and intracellular organization of translation across bacteria

**DOI:** 10.1101/479840

**Authors:** William T. Gray, Sander K. Govers, Yingjie Xiang, Bradley R. Parry, Manuel Campos, Sangjin Kim, Christine Jacobs-Wagner

## Abstract

The scaling of organelles with cell size is thought to be exclusive to eukaryotes. Here, we demonstrate that similar scaling relationships hold for the nucleoid in bacteria. Despite the absence of a nuclear membrane, nucleoid size strongly correlates with cell size, independent of changes in DNA amount and across various nutrient conditions. This correlation is observed in diverse bacteria, revealing a near-constant ratio between nucleoid and cell size for a given species. As in eukaryotes, the nucleocytoplasmic ratio in bacteria varies greatly among species. This spectrum of nucleocytoplasmic ratios is independent of genome size, and instead appears linked to the average cell size of the population. Bacteria with different nucleocytoplasmic ratios have different biophysical properties of the cytoplasm, impacting the mobility and localization of ribosomes. Together, our findings identify new organizational principles and biophysical features of bacterial cells, implicating the nucleocytoplasmic ratio and cell size as determinants of the intracellular organization of translation.

## Introduction

The spatial organization of the cell has a profound effect on various cellular processes from bacteria to humans (Bisson-Filho et al., 2018; Diekmann and Pereira-Leal, 2013; Harold, 2005; Surovtsev and Jacobs-Wagner, 2018). In eukaryotic cells, a distinctive feature of intracellular organization is the nucleus, a membrane-enclosed organelle that harbors most of the cell’s genetic material. The nuclear envelope hereby spatially confines the genetic material and physically separates transcription and translation. While the sizes of cells and nuclei vary considerably among species and tissues, there is a remarkable linear size scaling relationship between the cell and the nucleus for a given cell type, which was first reported over 100 years ago (Conklin, 1912; Woodruff, 1913). Correlations between cell size and nuclear size are not only widespread among eukaryotic cells but also robust to genetically-and nutritionally-induced cell size perturbations (Jorgensen et al., 2007; Neumann and Nurse, 2007). This scaling property results in a constant ratio between nuclear and cellular volumes, also known as the karyoplasmic or nucleocytoplasmic (NC) ratio (Wilson, 1925). Why cells maintain a specific NC ratio is generally not well understood, though alterations in NC ratios have been associated with aging and diseases such as cancer (Capell and Collins, 2006; Chow et al., 2012; Prokocimer et al., 2009; Zink et al., 2004). The sizes of other cellular components such as vacuoles, mitotic spindles, centrosomes and mitochondria have also been shown to scale with cell size in various eukaryotic cell types (Levy and Heald, 2012; Marshall, 2015; Reber and Goehring, 2015). As such, these scaling properties are believed to be unique to eukaryotes.

In bacteria, the chromosomal DNA typically occupies a subcellular region called the nucleoid (Kellenberger et al., 1958; Mason and Powelson, 1956). Recently, we showed that the average size of the nucleoid scales with the average size of the cell across ∼4,000 gene-deletion mutants of *Escherichia coli* (Campos et al., 2018). In addition, nucleoid size and cell size in *E. coli* correlate at the single-cell level, at least under specific growth conditions (Junier et al., 2014; Paintdakhi et al., 2016). An intuitive explanation for these observations may be linked to differences in DNA amount. Even under nutrient-poor conditions, DNA replication happens during a large part of the cell cycle, such that bigger cells tend to contain more DNA. This is exacerbated under nutrient-rich conditions under which *E. coli* displays overlapping DNA replication cycles (Cooper and Helmstetter, 1968). This leads to a continuous increase in DNA content from cell birth to division (Cooper and Helmstetter, 1968). Recent work with mutants of altered cell widths further suggests that the amount of DNA in such rapidly growing cells is directly coupled to cell volume (Shi et al., 2017). However, whether the scaling of nucleoid size with cell size is exclusively linked to changes in DNA content remains to be established. It is also currently unclear whether a scaling relationship between nucleoid and cell size is robust across growth conditions or widespread among bacteria. At the same time, it is unclear whether the size of the nucleoid or the volume fraction it occupies within the cell has any physiological consequence. We address all of these unknowns below.

## Results

### Nucleoid size scaling is robust across a wide range of cell sizes in *E. coli*

Given that different nutrient conditions give rise to cells of different sizes (Pierucci, 1978; Schaechter et al., 1958), we used phase contrast and fluorescence microscopy to examine how cell size variation in exponentially growing *E. coli* may affect nucleoid size across 30 nutrient conditions (M9 medium supplemented with different carbon sources ± casamino acids and thiamine, see Table S1). Cell contours were detected and curated in an automated fashion using the open source software package Oufti (Paintdakhi et al., 2016) and a support vector machine model (see STAR Methods). For each condition, the areas of thousands of cells were quantified from the cell contours (Figure 1A, Figure S1A). DAPI-stained nucleoids were detected using the objectDetection module of Oufti, from which we extracted the total area occupied by the DAPI signal (Figure 1A). Since estimation of the nucleoid area can vary with the chosen Oufti parameters (e.g., contour rigidity, relative signal threshold), we used the same parameter values across growth conditions.

**Figure 1.**
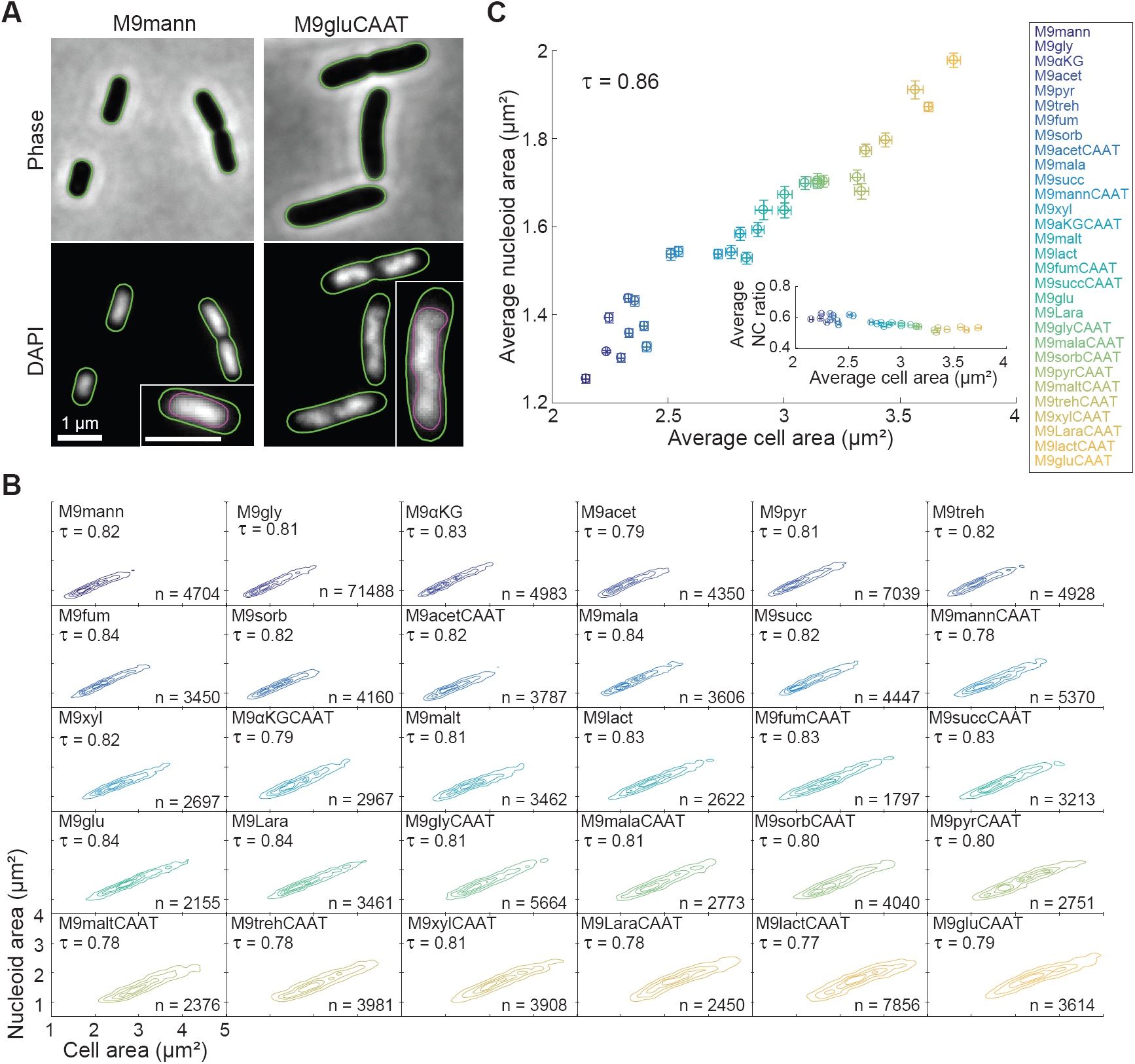
Nucleoid size scaling is robust across a wide range of *E. coli* cell sizes. A. Phase contrast and DAPI images of *E. coli* cells (CJW6324) grown in liquid cultures of M9 medium supplemented with 0.2% mannose (M9mann) or 0.2% glucose, 0.1% casamino acids and 1μg/ml thiamine (M9gluCAAT) at 37 °C. The images were processed with Oufti to identify the contours of the cells (green) and nucleoids (purple, insets). B. Density contour plots showing the strong correlation between cell area and nucleoid area for individual CJW6324 cells grown in the indicated growth media (for a full description of the growth media, see Table S1). The contour lines represent the 0.10, 0.25, 0.50 and 0.75 probability envelopes of the data. C. Scatter plot of the average cell area versus the average nucleoid area for the indicated growth conditions. Inset: scatter plot of the average cell area versus the average NC ratio for the same growth conditions. Error bars indicate 95% confidence intervals. See also Figures S1-3.

Using this methodology, we observed a strong correlation (Kendall correlation τ ≥ 0.77) between the cell area and nucleoid area of individual cells within all 30 tested growth conditions (Figure 1B). These results show that the nucleoid size scaling property is robust across a wide range of growth rates, with doubling times varying from ∼40 min to ∼4 h (Figure S1B). For each condition, the nucleocytoplasmic (NC) ratio (nucleoid area divided by cell area) was independent of the total or normalized intensity of the DAPI signal per cell (Figure S2), and was therefore unaffected by variations in DAPI staining efficiency. Moreover, we observed identical scaling relationships between nucleoid and cell area for nucleoids labeled with an mCherry or CFP fusion to a subunit of the nucleoid-associated HU complex (Figure S3A-C). The scaling between the cell area and the total nucleoid area was preserved in filamentous cells obtained by treatment with cephalexin (Figure S3D), a drug that inhibits cells division without affecting growth and DNA replication (Boye and Lobner-Olesen, 1991; Rolinson, 1980). The scaling relationship in these filamentous cells was almost indistinguishable from that in untreated cells (Figure S3D). These observations indicate that nucleoid size scaling occurs independently of cell division and persists across a wide range of cell sizes and growth conditions.

At the population level, we also observed a strong correlation (τ = 0.86) between the mean cell area and the mean nucleoid area of untreated cells across the tested 30 growth conditions (Figure 1C). This relationship was not perfectly linear, as the average NC ratio slightly decreased with increasing average cell size (Figure 1C, inset). This small decrease, which will be addressed later, was not a consequence of differences in growth medium osmolality (Figure S1C), which can cause variations in nucleoid morphology (Cagliero and Jin, 2013).

### Nucleoid size scaling is independent of DNA replication

We next investigated whether changes in DNA content underlie the scaling of nucleoid size with cell size by using nutrient-poor growth conditions. In such environments, *E. coli* cells display discrete cell cycle periods, known as the B, C, and D periods, corresponding to cell-cycle phases before, during, and after DNA replication, respectively (Cooper and Helmstetter, 1968). If DNA replication was solely responsible for nucleoid size scaling, we would expect to observe a correlation between nucleoid and cell size only during the C period, and not during the B and D periods when the DNA amount does not change. As cell size and the DAPI signal intensity did not provide sufficient resolution to distinguish between cells in the B, C, and D periods (Figure S4A), we used a strain producing a SeqA-mCherry fusion. SeqA associates with newly replicated DNA by transiently binding hemi-methylated GATC sites (Brendler et al., 1995; Lu et al., 1994; Slater et al., 1995). When fluorescently tagged, SeqA forms bright fluorescent foci that trail the replication forks during DNA replication (C period). In the absence of DNA replication (B and D periods), SeqA-mCherry displays diffuse nucleoid-associated fluorescence (Adiciptaningrum et al., 2015; Helgesen et al., 2015; Molina and Skarstad, 2004; Wallden et al., 2016) (Figure 2A). By quantifying the relative area of the SeqA-mCherry signal and combining this information with cell area measurements, we were able to identify three distinct groups of cells—corresponding to the B, C, and D cell cycle periods—in populations growing under various nutrient-poor conditions (Figure 2A and Figure S4B-D). Surprisingly, we found a strong correlation of nucleoid area with cell area for all three periods (Figure 2B and Figure S4D). The correlations and slopes were the strongest in the C period under all 11 tested nutrient-poor conditions, but both remained significant during the B and D periods (Figure 2B and Figure S4B-C). Apart from these small differences between cell cycle periods, we observed similar average NC ratios for each growth condition (Figure S4B). These results indicate that the scaling between nucleoid and cell sizes occurs independently of DNA replication.

**Figure 2.**
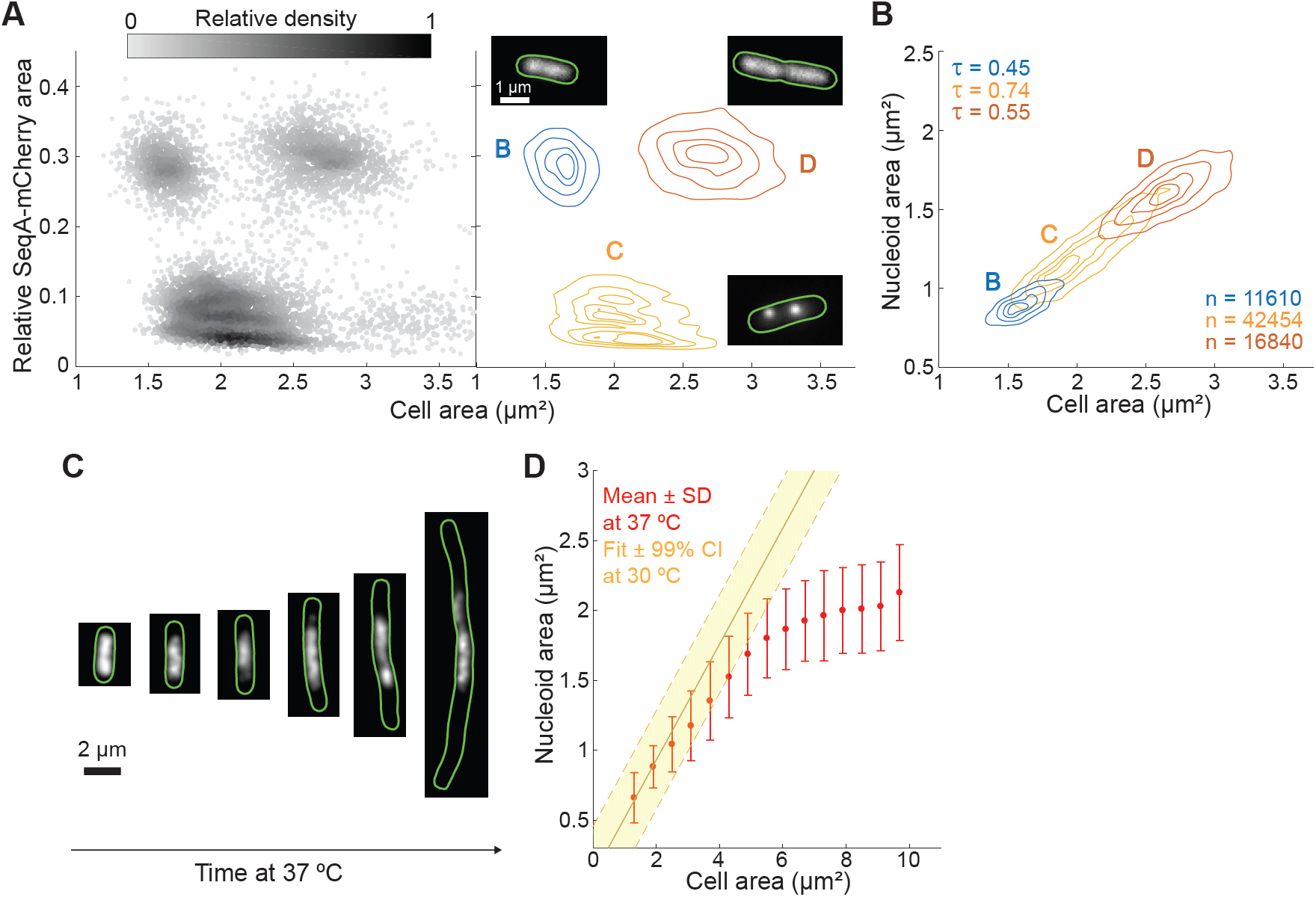
Nucleoid size scaling with cell size does not depend on DNA replication. A. Density scatter plot (left) and density contour plot (right) of cell area versus the relative SeqA-mCherry signal area of *E. coli* cells (CJW6324) grown in M9 medium supplemented with 0.2% glycerol. The gray scale in the density scatter plot indicates the relative density of dots (cells) in a given area of the chart. This plot was used to identify cells in the B, C and D cell cycle periods found under these growth conditions. The contour lines represent the 0.10, 0.25, 0.50 and 0.75 probability envelopes of the data. Insets: representative images of the subcellular SeqA-mCherry signal in a specific cell cycle period. B. Density contour plots of cell area versus nucleoid area for cells in B, C and D periods based on the analysis shown in A. The contour lines represent the 0.10, 0.25, 0.50 and 0.75 probability envelopes of the data. The nucleoid was detected by DAPI staining. See also Figure S4. C. Representative fluorescence images of *dnaC2* cells (CJW6370) producing HU-mCherry at different time points after a shift to a restrictive temperature (37 °C). D. Plot showing the average nucleoid area per cell area bin for HU-mCherry-labeled *dnaC2* cells at 37 °C. Cells (n = 12268) from different time points following temperature shifts were combined into one dataset and grouped into bins based on their cell areas. Shown are the average nucleoid area and standard deviation (SD) of each cell area bin. The solid yellow line indicates the expected relationship between nucleoid and cell area based on the scaling observed under permissive conditions (30 °C). The dotted lines indicate the 99% confidence interval (CI) of the fit.

To confirm this unexpected conclusion, we used temperature-sensitive *dnaC2* mutant cells producing an HU-mCherry fusion to visualize the nucleoids. At restrictive temperatures, these cells are unable to initiate new rounds of DNA replication, but continue to grow without dividing (Carl, 1970). We found that 90 min after the temperature shift to 37 °C, the average cell size of the population began increasing at which time we measured the size of both cells and nucleoids at regular intervals for 210 min. Remarkably, the size of the nucleoid increased with cell size over an almost 4-fold range before reaching a plateau in long cells (Figure 2C-D). Before reaching this limit, the scaling relationship in the absence of DNA replication was similar to that observed under the permissive temperature (30 °C) when DNA replication occurs (Figure 2C-D). Together, these observations demonstrate that nucleoid size scaling occurs irrespective of changes in DNA content.

### The nucleoid size scaling property is conserved in *Caulobacter crescentus*, but with a different NC ratio

To examine whether a scaling relationship between nucleoid and cell size is observed in other bacteria, we imaged DAPI-stained *C. crescentus* cells expressing CFP-labeled DnaN. DnaN is the β sliding clamp of the DNA polymerase, which, when fluorescently labeled, forms foci during DNA replication but otherwise displays a disperse distribution (Arias-Cartin et al., 2017; Collier and Shapiro, 2009; Fernandez-Fernandez et al., 2013). By quantifying the signal area of DnaN-CFP, we were able to readily identify cells in distinct cell-cycle periods (Figure 3A). As with *E. coli* (Figure 2B), we observed a strong scaling relationship between nucleoid size and cell size in cells in the B and D periods (Figure 3B), indicating that nucleoid size scaling occurs even in the absence of DNA replication. As in *E. coli*, nucleoid size determination in *C. crescentus* was independent of DAPI signal intensity (Figure S5A), and insensitive to the nucleoid labeling method (Figure S5B). Scaling was maintained in defined (M2G) and complex (PYE) growth media (Figure S5C) as well as in mutants with altered cell sizes and morphologies (Figure S5D-E), such as FtsZ-depleted, δ*rodZ* and δ*hfq* cells (Alyahya et al., 2009; Irnov et al., 2017; Wang et al., 2001).

**Figure 3.**
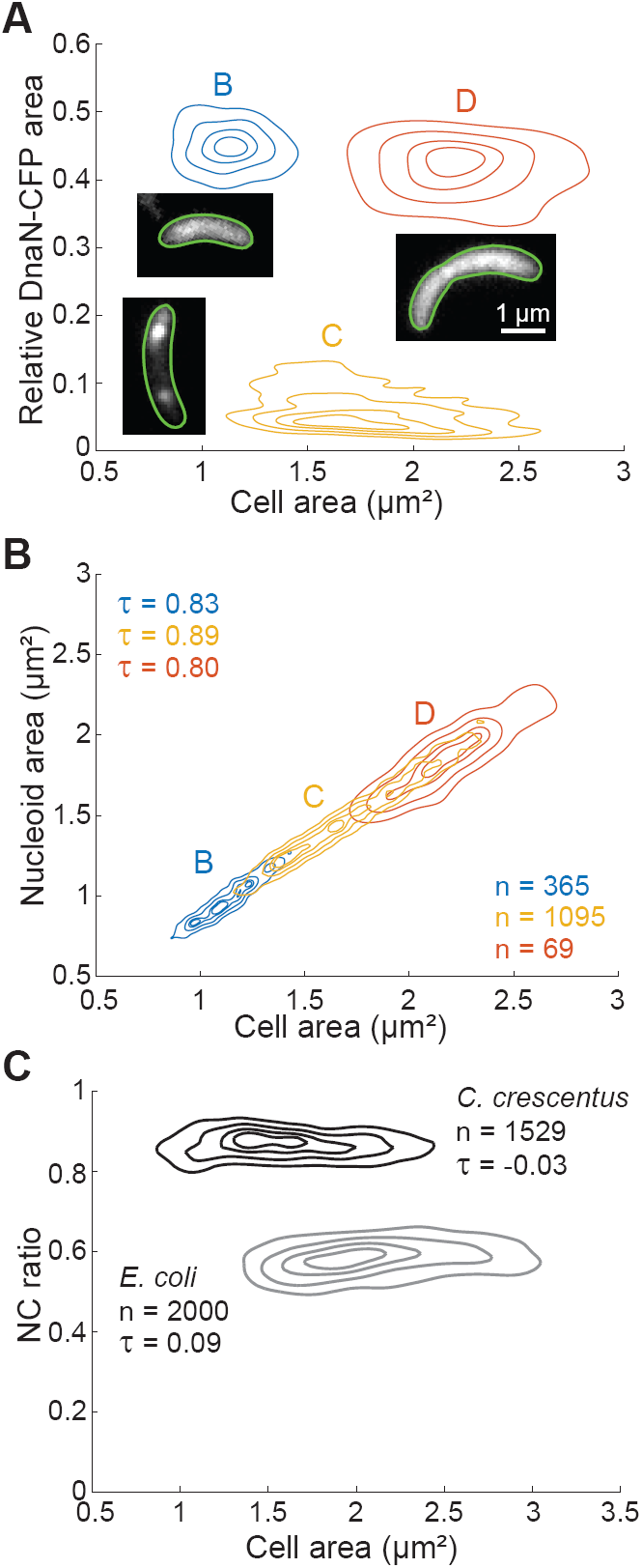
Nucleoid size scaling is also observed in *C. crescentus*, a bacterium with a different NC ratio. All contour lines represent the 0.10, 0.25, 0.50 and 0.75 probability envelopes of the data. A. Density contour plot of cell area versus the relative DnaN-CFP signal area of *C. crescentus* cells producing a DnaN-CFP fusion (CJW5969) and grown in M2 medium supplemented with 0.2% glucose. This plot was used to identify cells in the B, C, and D cell cycle periods. Insets: representative images of the subcellular DnaN-CFP signal in a specific cell cycle period. B. Density contour plots of cell area versus nucleoid area for cells in panel A. The nucleoid was detected by HU-mCherry labeling. C. Density contour plots of cell area versus NC ratio for *E. coli* (CJW6324) and *C. crescentus* (CJW5969) cells grown in M9 medium supplemented with 0.2% glycerol and M2 medium supplemented with 0.2% glucose, respectively. The nucleoid was detected by DAPI staining for *E. coli* and by HU-mCherry labeling for *C. crescentus*. See also Figure S5.

Nucleoid size scaled with cell size in both *E. coli* and *C. crescentus*. However, their NC ratios were very different (Figure 3C). This is consistent with observations that the nucleoid spreads through most of the cell in *C. crescentus* whereas *E. coli* displays DNA-free regions (Jensen and Shapiro, 1999; Kellenberger et al., 1958). The large NC ratio in *C. crescentus* was not due to PopZ-mediated attachment of the chromosome to the cell poles (Bowman et al., 2008; Ebersbach et al., 2008), as it was maintained in the Δ*popZ* mutant (Figure S5E-F).

### Nucleoid size scaling across bacterial phyla reveals a continuum of NC ratios

The scaling relationship between nucleoid and cell sizes is likely a common bacterial feature, as we observed it in over 35 species from different phyla or classes (Figure 4A and Figure S6A). Each species investigated displayed a constant, specific NC ratio (Figure 4B). To avoid measurement biases, we used the same Oufti parameters to identify the nucleoid contour of all cells in this dataset. As with *E. coli* and *C. crescentus*, we confirmed that the NC ratio was not affected by the intensity of the DNA signal (Figure S7A-B). We also observed no correlation between the average DNA signal intensity and the average NC ratio (Figure S7C). The various species were generally grown in complex media described in the literature or recommended by the provider. In some cases, we examined different growth conditions. For example, we imaged *Bacteroides thetaiotaomicron* (*B. theta*) grown *in vitro* in both complex (TYG) and defined (GMM) media, or *in vivo* in mono-associated gnotobiotic mice. For the latter, the samples were obtained from the cecum and feces. These different growth conditions revealed differences in cell sizes but, in all cases, nucleoid size scaled with cell size at the single-cell level (Figure 4A).

**Figure 4.**
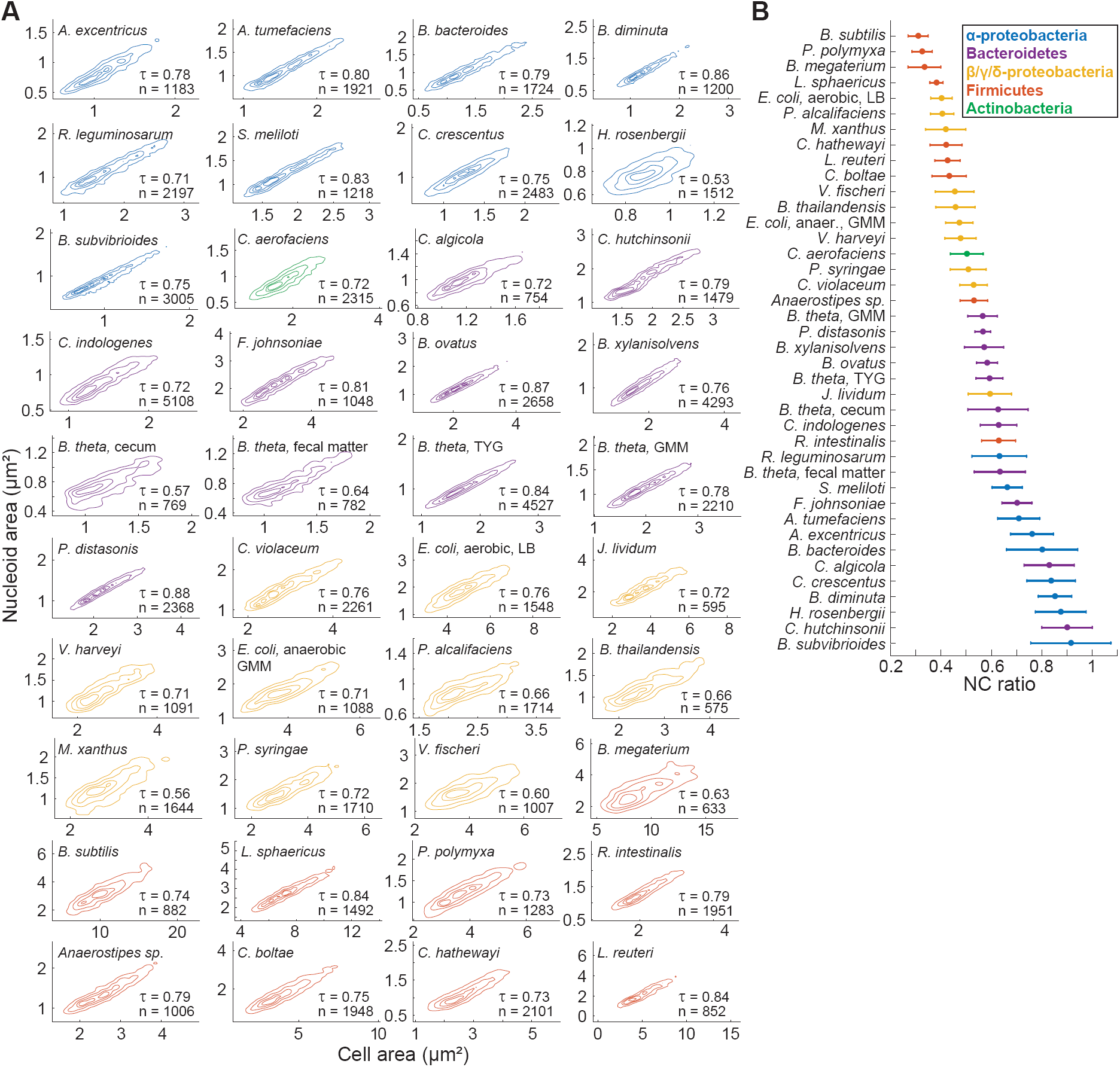
Nucleoid size scaling across bacterial species from different phyla reveals a continuum of NC ratios. A. Density contour plots of cell area versus nucleoid area for fixed cell populations from different bacterial species. The contour lines represent the 0.10, 0.25, 0.50 and 0.75 probability envelopes of the data. When different growth conditions were examined for the same species, the growth medium is indicated next to the species name. Contours of the same color indicate affiliation to the same phylum or class. The DNA dye used for nucleoid labeling for each species is detailed in the STAR methods. B. Average NC ratio (with error bars representing the standard deviation) for all the included species. See also Figures S6-7.

The name “nucleoid” (nucleus-like) comes from the early observation that the bacterial chromosome occupies a distinct intracellular region (Kellenberger et al., 1958; Mason and Powelson, 1956), as exemplified by the organization of the γ-proteobacterium *E. coli* (Figure 1A). The near-cell-filling organization of the chromosome in the α-proteobacterium *C. crescentus* is usually ignored or thought of as an exception (Campos and Jacobs-Wagner, 2013; Surovtsev and Jacobs-Wagner, 2018). Analysis of the average NC ratios of our panel of diverse species revealed that high average NC ratios, i.e., near-cell-filling nucleoids, can be found not only in other α- proteobacteria but also in some Bacteroidetes (Figure 4B). Furthermore, there was no subdivision of the analyzed bacteria into discrete lower and higher NC ratio categories. Instead, we observed a continuum of average NC ratios across species (Figure 4B).

While sorting species based on their average NC ratios revealed some phylogenetic clustering (Figure 4B), phylum association was not necessarily predictive of NC ratio. For example, α-proteobacteria generally had a higher NC ratio than proteobacteria from the β, γ, or δ classes (Figure 4B). Bacteroidetes provided a striking example of distinct chromosome organization within a phylum. *Cytophaga hutchinsonii* displayed a high NC ratio, characteristic of cell-filling DNA, whereas *Parabacteroides distasonis* exhibited a considerably lower NC ratio and clear DNA-free regions (Figure 4B and Figure S6A). These results indicate that the intracellular organization of the chromosome is an evolvable feature that varies significantly between species without strict phylogenetic determinants.

### The average NC ratio negatively correlates with the average cell size

Given this surprisingly large spectrum of average NC ratios among bacteria, we wondered whether certain cellular characteristics are associated with a given NC ratio. We found no correlation between genome size and average NC ratio (or average nucleoid area, or cell volume), despite a ∼3-fold difference in genome size between the included species (Figure S6B). Growth rate was also a poor predictor of NC ratios. Fast-growing species such as *E. coli* (in LB), *Bacillus subtilis* (in LB) and *B. theta* (in TYG medium), which have doubling times of ∼20 to ∼30 min at 37 °C (Eley et al., 1985; Taheri-Araghi et al., 2015; Weart et al., 2007) displayed a wide range of NC ratios, whereas the NC ratio of the slower-growing *Myxococcus xanthus* (in CYE medium), which has a doubling time of ∼4 h (Sun et al., 1999), was similar to that of *E. coli* growing in LB.

We did, however, observe a striking, seemingly exponential relationship between the average cell volume and the average NC ratio of bacteria (Figure 5A). The exponential relationship was particularly apparent upon plotting the mean NC ratio versus the logarithm of the mean cell volume (Figure 5A, inset), with a Kendall correlation τ = −0.70. This strong correlation indicates that the average cell volume of a species is highly predictive of the average NC ratio. We also observed strong relationships between other morphological descriptors and the NC ratio (Figure 5B). Further underscoring the validity of these relationships is the fact that the *E. coli* data from cultures grown under 30 different nutrient conditions (Figure 1C) overlapped almost perfectly with the curve obtained with the different bacterial species (Figure 5A, inset). The negative relationship between average NC ratio and average cell size observed in these experiments (Figure 1C) thus appears to be a consequence of the more general relationship between these two cellular characteristics.

**Figure 5.**
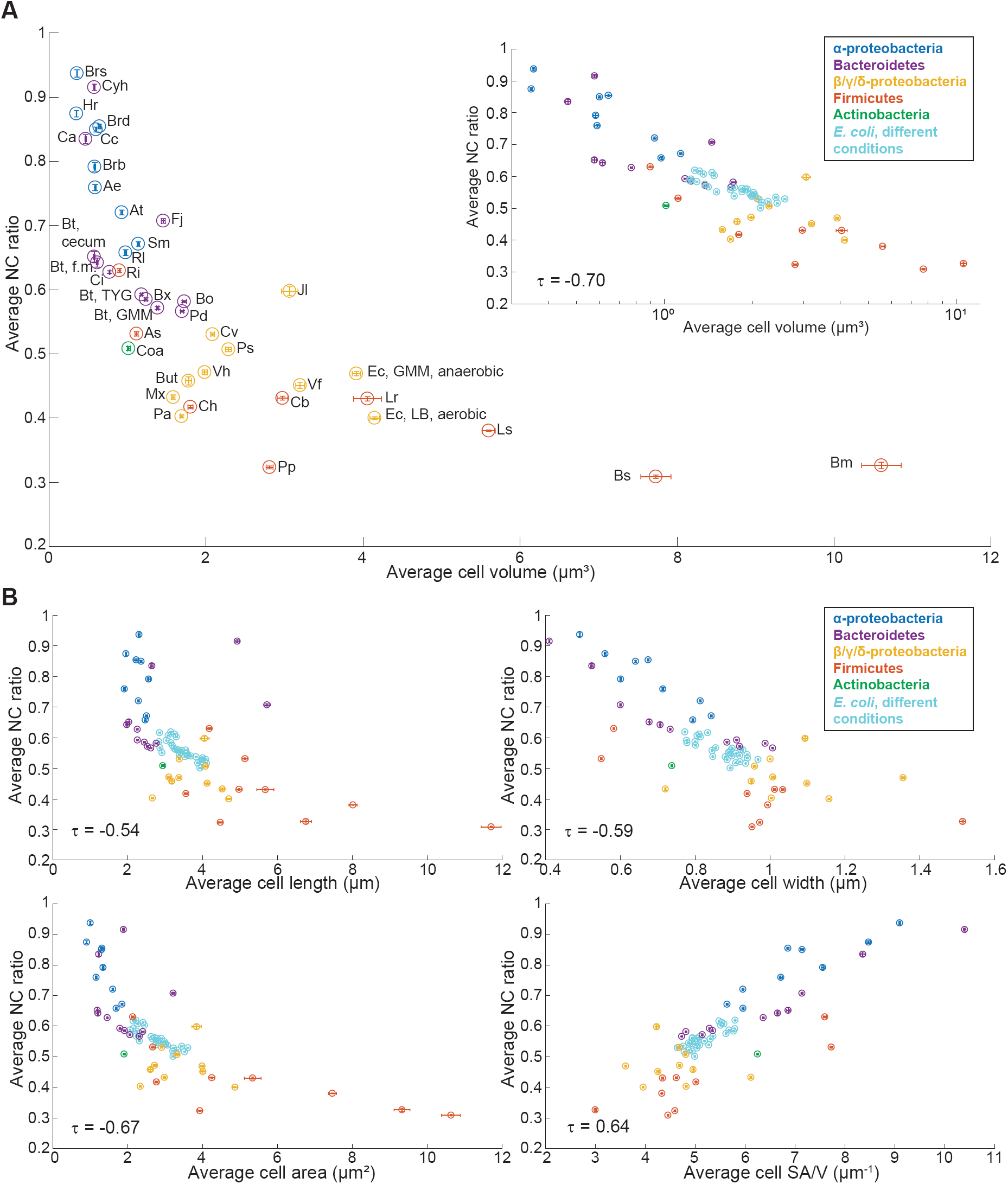
The average NC ratio is linked to the average cell size. For all plots, error bars indicate 95% confidence intervals. A. Scatter plot of average cell volume versus average NC ratio for all the included species. Abbreviated species names are indicated next to the corresponding datapoint; see Key Resources Table for a full name description. Inset: same relationship with average cell volume on a logarithmic scale. B. Scatter plot of average NC ratio versus average cell length, average cell width, average cell area or average surface area to volume ratio.

### The cytoplasm of bacteria with different NC ratios displays different biophysical properties

What are the physiological implications of a high or low NC ratio? We speculated that DNA might affect the dynamics, and thereby the organization, of large cellular components whose diffusion may be impeded by the DNA meshwork. In bacteria with low NC ratios like *E. coli*, large objects may be able to more freely diffuse in DNA-free regions. In contrast, motion may be limited in bacteria with high NC ratios like *C. crescentus* where the DNA spreads throughout most of the cytoplasm. To test this idea, we conducted experiments using genetically-encoded GFP-μNS particles expressed in *E. coli* and *C. crescentus*. We previously showed that GFP-μNS particles are useful to probe the biophysical properties of the bacterial cytoplasm (Parry et al., 2014). These probes derive from a mammalian reovirus protein that assembles into spherical objects (Broering et al., 2005; Broering et al., 2002). Once fused to GFP, they form fluorescent particles that increase in signal intensity and absolute size with increased GFP-μNS synthesis (Parry et al., 2014). We tracked GFP-μNS particles from three bins of particles of similar intensity (and, consequently, size) in both *E. coli* and *C. crescentus* growing at a similar rate (Figure 6A and Movie S1-2). Comparison of the ensemble-averaged mean squared displacements (MSDs) for particles belonging to these bins revealed drastic differences in probe dynamics between the two species (Figure 6B). GFP-μNS particles in *C. crescentus*, independent of their size range, displayed significantly lower mobility than in *E. coli* (Figure 6B). Diffusion measurements of free GFP are similar in these two species (Elowitz et al., 1999; Montero Llopis et al., 2012), indicating that a difference in cytoplasmic viscosity cannot explain these observations. Instead, these observations support the notion that different NC ratios can lead to different biophysical properties of the cytoplasm that affect the mobility of large cytoplasmic objects.

**Figure 6.**
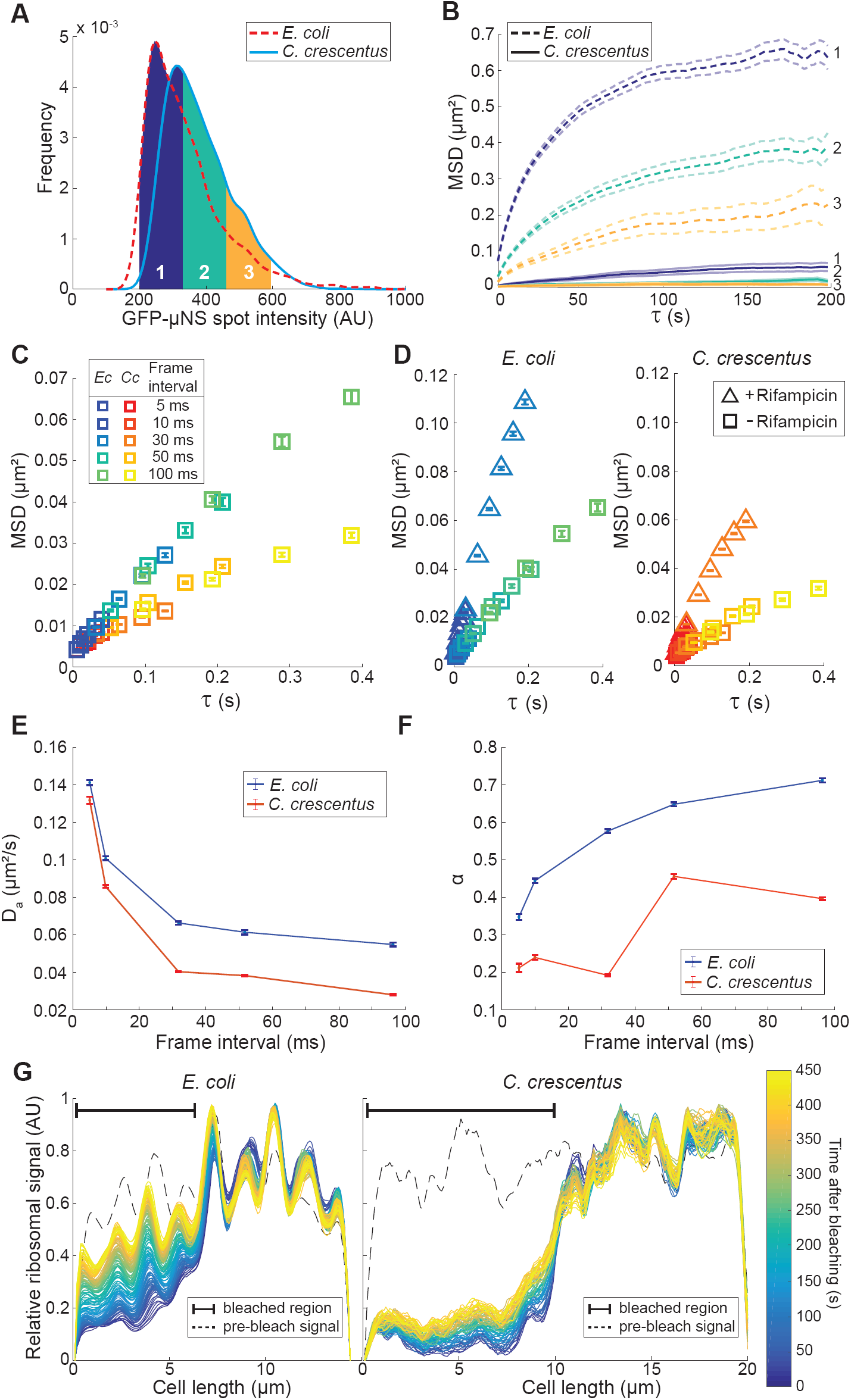
The intracellular mobility of large objects displays non-linear dynamics and is different between *E. coli* and *C. crescentus*. *E. coli* cells (CJW6723) were grown in M9 medium supplemented with 0.2% glycerol and *C. crescentus* cells (CJW6917) in M2 medium supplemented with 0.2% glucose, resulting in similar doubling times of ∼120 min. A. Frequency distributions of GFP-μNS spot intensities in *E. coli* (n= 2279) and *C. crescentus* (n = 2019) cells. Three bins of GFP-μNS particles with similar intensities and thus sizes are indicated in color. B. Ensemble-averaged MSDs of GFP-μNS particles (belonging to the intensity bins highlighted in panel A) in *E. coli* cells (n = 1208, 600 and 200 for bins 1, 2 and 3, respectively) and *C. crescentus* cells (n = 837, 984 and 374 for bins 1, 2 and 3, respectively). Error bars indicate 95% confidence intervals. C. Ensemble-averaged MSDs of fluorescently-labeled ribosomes in *E. coli* (SX289) and *C. crescentus* (CJW5156) at different acquisition frame intervals. For each frame interval, > 8900 trajectories were collected. Only the first four points of the MSDs are shown for each frame interval. Error bars indicate 95% confidence intervals. D. Same as panel C, except that ensemble-averaged MSDs from rifampicin-treated cells (200 μg/ml, 2 h) were added for comparison to the results shown in panel A. For each frame interval, > 3700 trajectories were collected in rifampicin-treated cells. For these MSDs, only the first six points are shown for the 5 ms and 30 ms frame intervals. The color scheme for the frame intervals is the same is in C. Error bars indicate 95% confidence intervals. E. Plot showing the apparent diffusion coefficients calculated from the aforementioned MSDs as a function of the frame interval. Error bars indicate 95% confidence intervals. F. Same as E, but for the anomalous exponent. G. Representative plots showing the evolution of the ribosomal fluorescence recovery over time (up to 450 s) along the length of a cephalexin-treated *E. coli* cell (CJW4677) and of an FtsZ-depleted *C. crescentus* cell (CJW3821) following photobleaching of about half of the cell. The dotted line shows the fluorescence profile prior to bleaching. See also Figure S8.

### Ribosome dynamics differ in bacteria with different NC ratio

What large cytoplasmic components may be impacted by differences in NC ratio? Under the conditions we used, GFP-μNS particles have reported sizes between 50 and 200 nm (Parry et al., 2014), a similar size range as polysomes (Brandt et al., 2009), which are mRNAs loaded with multiple ribosomes (Miller et al., 1970; Warner et al., 1962). If polysome mobility is impacted by the DNA meshwork and the fraction of cellular space it occupies, it may explain a currently unresolved discrepancy in mRNA localization in the literature. Fluorescence *in situ* hybridization (FISH) microscopy experiments on several mRNAs in *C. crescentus* suggest that these mRNAs remain close to their corresponding gene loci throughout most of their lifetime (Montero Llopis et al., 2010). In contrast, a genome-wide FISH study in *E. coli* reveals no spatial enrichment of mRNAs near the corresponding chromosomal regions (Moffitt et al., 2016). Because translation starts on nascent mRNAs, polysomes are expected to form within the nucleoid. However, in *E. coli*, the low NC ratio creates DNA-free regions in which polysomes can more freely diffuse once they escape the DNA meshwork, leading to their dispersion. Conversely, the high NC ratio of *C. crescentus* would prevent the escape of polysomes from the DNA meshwork.

To test this hypothesis, we used photoactivated localization microscopy to track ribosomes in both *E. coli* and *C. crescentus*. In *E. coli*, we labeled ribosomes using a fusion of ribosomal subunit protein S22 with mEos2 (Wang et al., 2011). In *C. crescentus*, we tracked L1-Dendra2-tagged ribosomes (Lim et al., 2014). In both cases, the fusion replaced the wild-type copy of the ribosomal gene at its native chromosomal locus (Lim et al., 2014; Wang et al., 2011). Importantly, we acquired data at five different frame intervals (between 5 and 100 ms) and constructed ensemble MSDs for each frame interval (> 8900 trajectories per frame interval, Figure 6C and Movie S3-4). We reasoned that polysomes diffusing in a DNA meshwork may experience caging and uncaging behaviors, as observed for probes diffusing in gels (Brangwynne et al., 2009; Cai et al., 2011; Guo et al., 2014; Wong et al., 2004). Tracking at multiple timescales may reveal such non-linear dynamics in MSDs. As the majority of ribosomes (∼75-80%) are engaged in translation in both organisms (Forchhammer and Lindahl, 1971; Lin et al., 2004; Montero Llopis et al., 2012; Phillips et al., 1969; Varricchio and Monier, 1971), most of our trajectories likely reflected polysome dynamics.

The MSDs indeed revealed non-linear dynamics, with polysomes in *C. crescentus* displaying lower mobility than those in *E. coli*, especially at the longer (subsecond) timescales (Figure 6C). The difference in MSDs was not due to polysomes “experiencing” cell membrane confinement sooner in *C. crescentus* because of its smaller size than *E. coli*, as higher MSD values were obtained in both organisms following treatment with the transcription initiation inhibitor rifampicin (Figure 6D). Rifampicin treatment results in mRNA depletion, thus converting all polysomes into smaller, and therefore faster, free ribosomes (Blundell and Wild, 1971) that explore more cellular space in the same amount of time (Figure 6D). This finding demonstrates that at the subsecond timescale, cell size does not limit polysome mobility in either organism, and that cell confinement is not responsible for the observed mobility difference between the two species.

The non-linear dynamics of ribosomes became particularly apparent when we calculated the apparent diffusion coefficient (D_a_) and the anomalous exponents (α) from the MSDs. The value for D_a_ is commonly extracted from the slope of the first few time lags of the MSD curve using the equation MSD = 4D_a_t (Michalet, 2010). Anomalous exponents were obtained from the slope of the first three points of the MSD vs. time (as anomalous diffusion in the cytoplasm is characterized by a power law scaling: MSD(t) ∝t^α^ (Bouchaud and Georges, 1990)). Generally, in biological studies, D_a_ and α are assumed to be constant over time, such that most single-molecule tracking experiments are done using only a single time frame. However, our analysis revealed a striking dependency of D_a_ and α on the timescales at which the measurements were made (Figure 6E-F). In both organisms, D_a_ decreased with longer timescales, while α increased. Furthermore, the difference in D_a_ between *E. coli* and *C. crescentus* increased with increasing time intervals (Figure 6E), and the α value was consistently lower for ribosomes in *C. crescentus* than in *E. coli* (Figure 6F). These differences indicate that ribosomes, the majority of which is contained within polysomes, are much more confined in the high NC ratio bacterium *C. crescentus* than in the low NC ratio bacterium *E. coli*.

In single-molecule tracking experiments, the frame rate is usually under 100 ms to ensure accurate localization determination. However, the lifetime of most bacterial mRNAs is on the minute timescale (Chen et al., 2015; Kristoffersen et al., 2012; Moffitt et al., 2016; Redon et al., 2005). Given the time-dependency of ribosome dynamics, we anticipated that the difference in spatial exploration of ribosomes between *E. coli* and *C. crescentus* would be even more apparent at the physiologically relevant timescale of minutes. This is indeed what we observed in fluorescence recovery after photobleaching (FRAP) microscopy experiments (Figure 6G). To minimize the effects of cell geometry and photobleaching location on the observed fluorescence recovery, we used filamentous cells that were unable to divide due to cephalexin treatment (*E. coli*) or FtsZ depletion (*C. crescentus*), as routinely done (Elowitz et al., 1999; Montero Llopis et al., 2012). In these filamentous cells, the NC ratio remained the same as in normal sized cells (Figure S3D and S5D-E). Ribosomes were labeled using a RplA-GFP fusion in both species. RplA is the 50S ribosomal subunit protein L1, previously used to examine ribosome localization in *B. subtilis* (Mascarenhas et al., 2001). Due to the heterogenous distribution of the ribosomal signal in *E. coli*, we were unable to quantify ribosome mobility with a simple one or two-state diffusion model, but we did observe clear qualitative differences in ribosomal recovery between the two species (Figure 6G, Movie S5-6). *E. coli* cells showed nearly complete fluorescence recovery at the photobleaching location after 450 s while *C. crescentus* cells often recovered less than 20% of their prebleached fluorescence intensity.

### Intracellular organization of translation is associated with the NC ratio and cell size

The decreased mobility of polysomes in *C. crescentus* is consistent with the notion that the cell-filling nucleoid impedes polysome motion in this species. In *E. coli*, on the other hand, polysomes display higher mobility likely because they can diffuse more freely and accumulate in DNA-free regions once they escape the DNA meshwork. This raises the intriguing possibility that the difference in NC ratio and its impact on ribosome mobility contribute to the striking difference in spatial organization of ribosomes and thus translation between these two organisms. In *E. coli*, as in other bacteria with low NC ratios like *B. subtilis* and *Lactococcus lactis*, ribosomes are enriched in the nucleoid-free regions of the cytoplasm (Azam et al., 2000; Bakshi et al., 2012; Lewis et al., 2000; Robinow and Kellenberger, 1994; van Gijtenbeek et al., 2016), resulting in partial segregation of transcription and translation. In *C. crescentus* and *Sinorhizobium meliloti*, two bacteria with high NC ratios, a large physical separation of ribosomes and DNA is not observed, as both are found throughout the cytoplasm (Bayas et al., 2018; Montero Llopis et al., 2010).

If the NC ratio does indeed affect the spatial organization of translation, we may expect to already see changes in ribosome localization in *E. coli* cells grown in different nutritional environments that lead to small variations in NC ratios (Figure 1C, inset). To test this expectation, we used an *E. coli* strain carrying a mEos2 fusion to a ribosomal protein (Sanamrad et al., 2014) and grew this strain under 12 growth conditions that result in slightly varying NC ratios. Although nucleoid exclusion of ribosomes was observed for each growth condition, the exclusion was more pronounced in cells with smaller average NC ratios. This is exemplified in Figure 7A showing a comparison between cells in a nutrient-rich medium (M9gluCAAT, average NC ratio = 0.53) and cells in nutrient-poor medium (M9gly, average NC ratio = 0.58). We quantified the average nucleoid exclusion of ribosomes by calculating the signal correlation factor (SCF), a metric that measures the correlation between two fluorescent signals (see STARS Method). An SCF of 1, 0 and −1 indicates that the two signals display perfect co-localization, independent localization and exclusion, respectively. We restricted the calculation of SCF to a specific “correlation area” within individual cells (Figure S8A) to minimize the effects of cell size and geometry on the correlation (see STAR methods). This quantification across 12 growth media with varying NC ratios confirmed the gradual increase in ribosome exclusion with decreasing NC ratio (increasing average cell size), as evidenced by the more negative average SCF values (Figure 7B).

**Figure 7.**
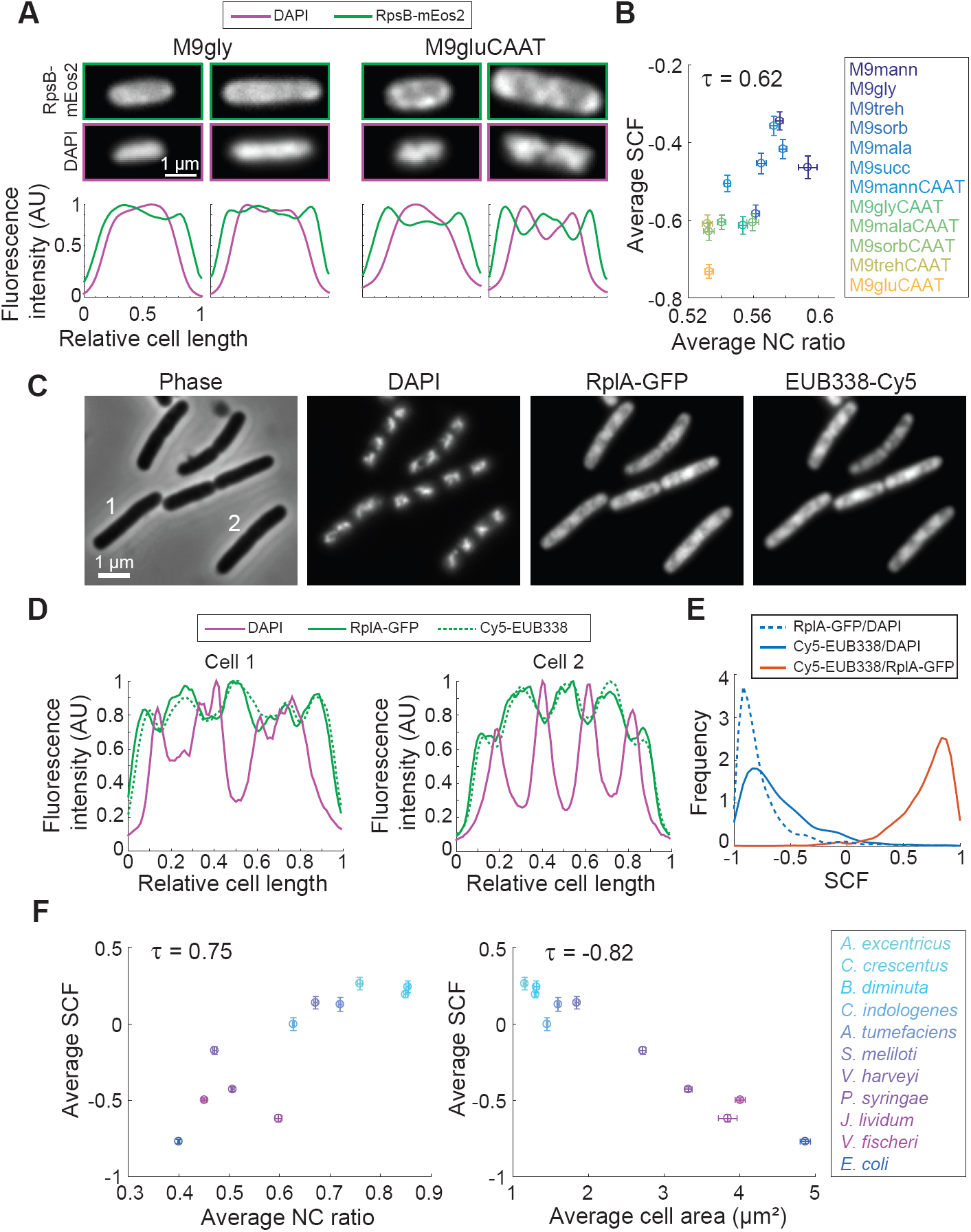
The spatial organization of ribosomes in bacteria is linked to the average NC ratio and cell size. A. Top, representative fluorescence images of *E. coli* cells (CJW6769) grown in M9 medium supplemented with 0.2% glycerol (M9gly) or 0.2% glucose, 0.1% casamino acids and 1μg/ml thiamine (M9gluCAAT). Bottom, fluorescence intensity profiles of DAPI and RpsB-mEos2 signals for these cells. B. Scatter plot of average SCF versus average NC ratio for *E. coli* cells (CJW6769) grown in the indicated growth media (for a full description of the growth media, see Table S1). The SCF was calculated by comparing the correlation between the DAPI and the RpsB-mEos2 signals for the indicated species. Error bars indicate 95% confidence intervals. C. Representative phase contrast and fluorescence images of *E. coli* cells (CJW4677) after the FISH procedure, highlighting the correspondence between the use of RplA-GFP and FISH (targeting 16S ribosomal RNA with the Cy5-labeled EUB338 probe) for visualizing ribosome localization. Cells were grown in M9 medium supplemented with 0.2% glycerol, 0.1% casamino acids, and 1μg/ml thiamine. D. Fluorescence intensity profiles of DAPI, RplA-GFP, and rRNA FISH (Cy5-EUB338) signals of *E. coli* cells (CJW4677) indicated in C, grown in M9 medium supplemented with 0.2% glycerol, 0.1% casamino acids, and 1μg/ml thiamine. E. Frequency distributions of SCF values between the rRNA FISH (Cy5-EUB338), RplA-GFP, and DAPI signals. F. Scatter plots of average SCF versus average NC ratio (left) or versus average cell area (right). The SCF was calculated by comparing the correlation between the DAPI and the rRNA FISH (Cy5-EUB338) signals for the indicated species. Error bars indicate 95% confidence intervals. See also Figure S8.

Given the continuum of NC ratios among diverse species (Figure 4B), we may also expect to see differences in ribosome localization among species with varying NC ratios. To examine this possibility, we performed fluorescence *in situ* hybridization (FISH) microscopy on 10 different species using a Cy5-labeled EUB338 probe complementary to the 5’ domain of 16S rRNA (Amann et al., 1990). This probe is complementary to the majority of eubacterial species sequenced and provides a method to visualize bulk ribosome localization in diverse species. As a control, we first performed SCF quantification for an *E. coli* strain producing fluorescently labeled ribosomes. This test revealed that cell fixation, a necessary step of the FISH procedure, slightly affects ribosome and DNA localization, thereby artificially increasing the SCF value (Figure S8B). Despite this caveat, we still observed nucleoid exclusion of ribosomes and strong colocalization between the ribosome signals obtained from the fluorescent labeling (using RlpA-GFP) and the FISH procedure (using Cy5-EUB338) at the single-cell and population levels (Figure 7D-E), validating our FISH method. For the 10 species tested, we found that the SCF obtained by FISH correlates with their average NC ratio (Figure 7F). Given the spectrum of NC ratios among diverse species (Figure 4B) and the correlation between the average NC ratio and the average cell size (Figure 5), our results also suggest a relationship between nucleoid exclusion of ribosomes and average cell size. Indeed, we found a strong negative correlation (τ = −0.82) between the average SCF and the average cell size across the tested species (Figure 7G). In other words, the bigger the average size of the species, the smaller its average NC ratio and the more ribosomes were excluded from the nucleoid. Altogether, our findings suggest a continuum of ribosome organization across bacteria and identify the average NC ratio and cell size of a species in a given growth medium as good predictors of how this bacterium spatially organizes translation.

## Discussion

Although the first reports of scaling relationships in eukaryotes between the size of subcellular components and that of the cell date back more than 100 years (Conklin, 1912; Marshall, 2015; Wilson, 1925; Woodruff, 1913), this phenomenon has remained largely unexplored in bacteria. Here, we demonstrate that nucleoid size strongly scales with cell size in exponentially growing cultures across a wide range of cell sizes and a diverse panel of bacterial species (Figures 1, 2, 3, 4, S1, S3D, S4, S5 and S6). Despite the apparently conserved nature of nucleoid size scaling, we found a continuum of NC ratios across species (Figure 4B), which can be predicted from the average cell size of the bacterial population (Figure 5). We highlight important biological implications of having a different NC ratio for the mobility and localization of larger particles such as polysomes (Figure 6), thereby implicating the NC ratio as an important determinant of the intracellular organization of bacterial translation (Figure 7).

Using the model bacteria *E. coli* and *C. crescentus*, we show that the scaling of nucleoid size with cell size occurs in the absence of changes in DNA content (Figure 2B-D and 3B). This is in line with findings in yeast cells in which increases in the amount of DNA do not directly lead to increases in nuclear size (Jorgensen et al., 2007; Neumann and Nurse, 2007). In eukaryotes, nuclear structural components, nucleocytoplasmic transport and nuclear envelope expansion have all been implicated in regulating nuclear size (Hara and Merten, 2015; Jevtic et al., 2014; Kume et al., 2017; Levy and Heald, 2010). The fact that the scaling property extends to bacteria, which lack a nuclear envelope, makes it even more remarkable. It highlights an intrinsic property of the DNA and the cell that predates the development of membrane-enclosed organelles such as the nucleus. The fact that it arises regardless of the way the genome is packaged into the cell (independently of nuclear membrane or histones) suggests that it is an ancient and basic cellular feature.

Although nucleoid size scaling is widespread among bacteria, the resulting NC ratios vary considerably (Figure 4B). Here again, this is similar to what is observed in eukaryotes where the NC ratio varies greatly among cell types (Ganguly et al., 2016; Jevtic and Levy, 2015; Jorgensen et al., 2007; Kume et al., 2017; Neumann and Nurse, 2007; Novakova et al., 2016; Su Lim et al., 2015). We found no link between NC ratio and chromosome size or growth rate of a given species (Figure S6). Instead, we discovered a remarkable relationship between the average cell size of a population and its average NC ratio, as the latter strongly correlated with morphological metrics that reflect average cell size (i.e., average cell volume, length, width, area and surface area to volume ratio) (Figure 5). Although the relationship is strongest for the average cell volume, the strong correlations with other size-related variables currently preclude us from associating the NC ratio with a specific morphological feature. It is important to note that while the relationship between average cell size and average NC ratio has predictive value at the population level, it does not extend to the single-cell level. This is evident from the maintenance of the NC ratio over the course of a cell cycle (Figure 1) and is further exemplified by the fact that an overlap in cell size between *C. crescentus* and *E. coli* does not lead to an overlap in NC ratio at the single-cell level (Figure 3C). These findings indicate that although a general relationship between average cell size and the NC ratio exists, the latter is controlled by factors other than cell size at the single-cell level.

Differences in NC ratio among species and across growth conditions (Figures 1C and 4B) have physiological implications. By comparing the motion of ribosomes in *E. coli* and *C. crescentus*, we found that their mobility is significantly decreased in cells with high NC ratios. Given that cytoplasmic viscosity is similar in *E. coli* and *C. crescentus* based on GFP diffusion measurements (Elowitz et al., 1999; Montero Llopis et al., 2012), this reduction likely arises because the diffusion of polysomes is impeded by the DNA meshwork. This difference was most pronounced on longer timescales due to the time-dependent properties of ribosome movement (both in terms of D_a_ and α). These non-linear dynamics of polysomes inside the bacterial cytoplasm reveal that the DNA affects the biophysical properties of the bacterial cytoplasm. Our data suggest that polysomes and other similarly sized objects experience local caging when they encounter the DNA mesh. An implication of such non-linear dynamics is that direct comparisons of diffusion coefficients without considering physiologically relevant timescales and differences in α can be misleading. For example, at short frame rates, polysomes may not diffuse far enough to be “aware” that they are within a DNA meshwork. As a result, the D_a_ values in *E. coli* and *C. crescentus* are relatively close to each other, consistent with previous determinations (Bakshi et al., 2012; Bayas et al., 2018; Sanamrad et al., 2014). This could lead to the interpretation that ribosome dynamics are the same in these organisms. We show that this is true only at the millisecond timescale, a timescale at which polysomes primarily experience protein crowding, which is similar in the two species (Elowitz et al., 1999; Montero Llopis et al., 2012). As the timescale increases, the D_a_ values decrease as polysomes increasingly experience the DNA mesh. This highlights the non-linear biophysical properties of the bacterial cytoplasm and stresses the importance of making diffusion measurements at different length and time scales.

The decrease in D_a_ values over time is most dramatic in *C. crescentus* (Figure 5E) where, unlike in *E. coli*, polysomes cannot escape the DNA meshwork because it fills most of the cell. By themselves, the differences in ribosome mobility in *E. coli* and *C. crescentus* could be attributed to other factors (e.g., fraction of nascent vs. mature mRNAs) than a difference in NC ratio between the two species. However, the decreased mobility of genetically encoded GFP-μNS particles in *C. crescentus* (in comparison to *E. coli*) supports our interpretation that different NC ratios give rise to different physical properties of the cytoplasm and have widespread implications for larger cellular components and their associated processes. The reduction of polysome mobility in *C. crescentus* explains why mRNAs remain in close proximity to their corresponding gene loci in this organism (Montero Llopis et al., 2010). In *E. coli*, on the other hand, polysomes would be able to escape the nucleoid due to the lower NC ratio, after which their increased mobility would lead to a more dispersed mRNA localization, as recently shown (Moffitt et al., 2016). Based on this interpretation, we anticipate that the NC ratio of a given bacterium, together with the lifetime of the mRNA, will dictate whether protein synthesis from this mRNA primarily occurs near the gene locus where the mRNA was transcribed, or away from it.

In eukaryotes, the term cytosol is used to designate the part of the cytoplasm that is not held by organelles. We propose that a similar distinction can be made in bacteria. Even without a membrane-enforced separation, the nucleoid (organelle) provides a distinct biophysical environment from the DNA-free region of the cytoplasm (cytosol). The spectrum of NC ratios across species and growth conditions suggests that the cytosolic fraction of a bacterial cell is far from fixed, and is instead an evolvable feature (Figure 4B). Although the NC ratio depends on the growth conditions for a given species, the actual fluctuations between conditions are small in comparison to the entire spectrum observed across species (Figures 1C and 5B). This observation may reflect unappreciated evolutionary constraints on intracellular organization and cell size for a given bacterial species.

## Supporting information

## Acknowledgements

We thank Drs. Nora Ausmees, Jacques Batut, Steven Lindow, Savithramma Dinesh-Kumar, Jeanne S. Poindexter, Jo Handelsman, Bonnie Bassler, Peter Greenberg, D_a_vid Zusman, Wade Winkler, Pamela Brown, Eric Stabb, Mark McBride, Sunny Xie, Andrew Goodman, as well as the ATCC Bacteriology collection and the Yale *E. coli* Genetic Stock Center, for providing strains used in this work. We would specifically like to thank Drs. Andrew Goodman and Bentley Lim for sharing fixed *B. theta* cells harvested from monocolonized mice, and Dr. Michael Zimmerman for providing strains and for his help with setting up anaerobic growth experiments. We also thank the Jacobs-Wagner laboratory for fruitful discussions and for critical reading of the manuscript. This work was partly supported by the National Institutes of Health (R01 GM065835 to C.J.-W.). S.K.G. was partly funded by a fellowship from the Belgian American Education Foundation (B.A.E.F.). C.J.-W. is an investigator of the Howard Hughes Medical Institute.

## Author contributions

Conceptualization, W.T.G., S.K.G. and C.J.-W.; Methodology, W.T.G., S.K.G., Y.X., B.R.P., M.C. S.K. and C.J.-W. Software, W.T.G., S.K.G., Y.X., B.R.P. and M.C.; Formal Analysis, W.T.G., S.K.G., Y.X. and B.R.P.; Investigation, W.T.G. and S.K.G.; D_a_ta Curation, W.T.G., S.K.G., Y.X. and B.R.P.; Writing – Original Draft, S.K.G. and C.J.-W.; Writing – Review & Editing, W.T.G., S.K.G., Y.X., B.R.P., M.C., S.K., and C.J.-W; Visualization, W.T.G. and C.J.-W.; Supervision, C.J.-W.; Project Administration, C.J.-W.; Funding Acquisition, C.J.-W.

## STAR Methods

### CONTACT FOR REAGENT AND RESOURCE SHARING

Further information and requests for resources and reagents should be directed to and will be fulfilled by the Lead Contact, Christine Jacobs-Wagner.

### EXPERIMENTAL MODEL AND SUBJECT DETAILS

#### Bacterial strains and growth conditions

Construction of strains and plasmids is detailed in Table S2.

To obtain steady-state growth conditions, cells were first inoculated in the appropriate growth medium and grown to stationary phase in culture tubes. Cells were subsequently re-inoculated into fresh medium by diluting them 1/10000 or more, and grown until they reached an optical density at 600 nm (OD_600_) of 0.1-0.3 (depending on the growth medium and organism) before sampling for microscopy.

*E. coli* cells were grown in LB medium (30 °C), gut microbiota medium (GMM; 30 °C) (Goodman et al., 2011) or M9 medium (37 °C) supplemented with 0.2% carbon source and, in certain instances, with 0.1% casamino acids and 1 μg/ml thiamine (CAAT). *C. crescentus* cells were grown at 30 °C either in PYE medium or M2G medium. *Sinorhizobium meliloti* (30 °C), *Psuedomonas syringae* (30 °C), *Janthinobacterium lividum* (25 °C), and *Burkholderia thailandensis* (30 °C) were grown in LB medium. *Rhizobium leguminosarum*, *Agrobacterium tumefaciens*, *Asticcacaulis excentricus*, *Chryseobacterium indologenes*, *Brevundimonas subvibrioides*, *Brevundimonas bacteroides*, and *Brevundimonas diminuta* were grown at 30 °C in PYE medium. *Vibrio harveyi* and *Vibrio fischeri* were grown at 30 °C in LBS medium. *Myxococcus xanthus* and *Flavobacterium johnsoniae* were grown at 30 °C in CYE medium. *Hirschia rosenbergii* was grown at 30 °C in marine broth medium (Difco, Fisher Scientific). *Cellulophaga algicola* was grown at 30 °C in DSMZ Medium 172. *Cytophaga hutchinsonii* was grown at 25 °C in CYE medium supplemented with 1% glucose. *Bacillus subtilis*, *Bacillus megaterium*, *Lysinibacillus sphaericus*, and *Paenibacillus polymyxa* were grown at 30 °C in nutrient broth medium. *Bacteroides ovatus*, *Bacteroides thetaiotaomicron*, *Bacteroides xylanisolvens*, *Parabacteroides distasonis*, *Chromobacterium violaceum*, *Providencia alcalifaciens*, *Roseburia intesinalis*, *Anaerostipes sp*., *Clostridium boltae*, *Clostridium hathewayi*, *Lactoacillus reuteri*, and *Collinsella aerofaciens* were grown at 37 °C in GMM (Goodman et al., 2011). *B. theta* was also grown at 37 °C in TYG medium (Bacic and Smith, 2008). Fixed *B. theta* cells isolated from the cecum and fecal matter of monocultured mice were a kind gift of the Andrew Goodman laboratory (Yale University). All cells that were grown in GMM or TYG medium were cultured anaerobically. The exact composition of all growth media is detailed in Table S1.

Cephalexin treatment of *E. coli* cells was performed by first growing the cells in the indicated growth medium as described above. Steady-state cultures were subsequently exposed to cephalexin (50 μg/ml) for a period of time corresponding to about two doublings of an unexposed population (1 to 6 h, depending on the growth medium) and then imaged.

For FtsZ depletion in *C. crescentus*, CJW3821 cells carrying *ftsZ* under the xylose-inducible promoter were grown to an OD_660_ of ∼0.1 at 30 °C in PYE medium containing 0.3% xylose for proper FtsZ synthesis. Cells were then spun down (5000 × g for 5 min) and washed with fresh PYE containing no xylose. FtsZ depletion was then performed by growing cells in PYE at 30 °C for 3-6 h.

## METHOD DETAILS

### Microscopy

Unless otherwise indicated, cells were imaged on agarose (1%) pads supplemented with the appropriate growth medium. For most experiments live cells were used, except for Figures 4 and 5 for which cells were first fixed with 4% formaldehyde and for Figure 7C-F for which cells were fixed and permeabilized for FISH microscopy (see below).

Phase contrast and epifluorescence imaging was performed on a Nikon Ti-E microscope equipped with a 100X Plan Apo 1.45 NA phase contrast oil objective (Carl Zeiss), an Orca-Flash4.0 V2 142 CMOS camera (Hamamatsu), and a Spectra X light engine (Lumencor). The microscope was controlled by the Nikon Elements software. The following Chroma filter sets were used to acquire fluorescence images: DAPI (excitation ET350/50x, dichroic T400lp, emission ET460/50m), CFP (excitation ET436/20x, dichroic T455lp, emission ET480/40m), GFP (excitation ET470/40x, dichroic T495lpxr, emission ET525/50m), YFP (excitation ET500/20x, dichroic T515lp, emission ET535/30m), mCherry/TexasRed (excitation ET560/40x, dichroic T585lpxr, emission ET630/75m) and Cy5.5 (excitation ET650/45x, dichroic T685lpxr, emission ET720/60m). Specialized microscopy setups used for FRAP experiments and single-molecule or single-particle tracking are detailed below.

### GFP-μNS experiments

For GFP-μNS experiments in *E. coli*, we used a published protocol (Parry et al., 2014). Briefly, *E. coli* strain CJW6723 was grown at 30 °C in M9 medium supplemented with 0.2% glycerol to an OD_600_ = 0.05-0.1. The synthesis of GFP-μNS was induced by the addition of 200-500 μM IPTG for 60-120 min. After induction, cells were spun down (5000 × g for 5 min) and washed with fresh M9 glycerol medium and grown for at least 60 min to allow for GFP maturation. For experiments in *C. crescentus*, strain CJW6723 was grown at 30 °C in M2 medium supplemented with 0.2% glucose to an OD_660_ = 0.05-0.1. GFP-μNS synthesis was induced by the addition of 0.3% xylose to the medium for 30-120 min. After induction, cells were spun down and washed with fresh M2G medium and grown for at least 60 min to allow for GFP maturation. Cells were then spotted on 1.5% agarose pads containing M9 glycerol (*E. coli*) or M2G (*C. crescentus*) and imaged every 2 s at 30 °C.

### Photoactivated localization and single-molecule tracking experiments

For photoactivated localization microscopy and single-molecule (ribosome) tracking, cover slips and glass slides were washed in the following manner: sonication in 1 M KOH (15 min), sonication in milliQ H2O (15 min) and sonication in 70% ethanol (15 min) with 3-5 milliQ H2O rinses between wash solution changes. Cleaned glass slides and cover slips were then dried with pressured air. Cells were spotted on a 1.5% agarose pad made with M9 medium supplemented with 0.2% glycerol, 0.1% casamino acids, and 1 μg/ml thiamine for *E. coli* or M2 medium supplemented with 0.2% glucose for *C. crescentus*. Imaging was performed with an objective heat ring set at 30 °C. All images were acquired on an N-STORM microscope (Nikon) equipped with a CFI Apo TIRF 100× oil immersion objective (NA 1.49), lasers (Agilent Technologies) emitting at 405 nm (0.1-10%) and 561 nm (15-100%), and a built-in Perfect Focus system. Raw single-molecule data were taken at a frame rate of 200 to 10 frames per second in a field of view of 36 × 36, 120 × 120, or 200 × 200 pixels (depending on the frame rate) with an Andor iXon X3 DU 897 EM-CCD camera (Andor Technology). Rifampicin treatment was performed by exposing cells to 200 μg/ml (*E. coli*) or 50 μg/ml (*C. crescentus*) rifampicin for 2 h in liquid culture before sampling and imaging.

### Fluorescence recovery after photobleaching experiments

For the FRAP experiments, filamentous cells (generated either by a 2 h treatment with 50 μg/ml cephalexin for *E. coli* or a 3-6 h FtsZ depletion in *C. crescentus*) were spotted on 1.5% agarose pads with M9 medium supplemented with 0.2% glycerol, 0.1% casamino acids, and 1 μg/ml thiamine or PYE. Cells were imaged at room temperature (∼22 °C) with a Nikon E80i microscope equipped with 100X Plan Apo 1.45 NA phase contrast objective and an Andor iXonEM+ DU-897 camera controlled by the Metamorph software. Fluorescence photobleaching was performed using a Photonic Instrument Micropoint laser system at 488 nm. Cells were imaged once before photobleaching, then bleached (for ∼0.5 s), and subsequently imaged at equal intervals (3-6 s for 450 s depending on whether *E. coli* or *C. crescentus* was imaged).

### Fluorescence in situ hybridization experiments

For FISH experiments, *E. coli* cells were grown in LB medium at 30 °C, *C. crescentus* cells were grown in PYE medium at 30 °C, and the other bacterial species were grown as described above. FISH was performed similarly to previous methods described by our laboratory (Kim and Jacobs-Wagner, 2018; Montero Llopis et al., 2010). Briefly, exponentially growing cells (OD_600_ < 0.3) were fixed in a 4% formaldehyde solution (4% formaldehyde, 30 mM NaHPO3 pH 7.5) for 15 min at room temperature followed by 30 min on ice. The samples were spun down (5000 × g for 3 min) and washed in phosphate-buffered saline (PBS) treated with diethyl pyrocarbonate (DEPC) 3 times. The cell pellets were resuspended in DEPC-treated PBS (8.0 g/l NaCl, 0.2 g/l KCl, 1.44 g/l Na_2_HPO_4_ and 0.24 g/l KH2PO_4_) and adhered to poly-L-lysine-coated coverslips. Cells were then lysed with 70% ethanol for 5 min at room temperature. Pre-hybridization was then performed with a 40% formamide, 2x saline-sodium citrate solution (SSC, 300 mM NaCl, 30 mM sodium citrate, pH 7.0) containing 0.2 mM vanadyl ribonucleoside complex (VRC) for 2 h at 37 °C. Immediately afterwards, hybridization was performed with EUB338 (5’-GCTGCCTCCCGTAGGAGT-3’, 5’-monolabeled with Cy5) in a solution containing 4 nM EUB338, 40% formamide, 2x SSC, 0.2 mM VRC, 10% dextran sulfate, 0.1% bovine serum albumin, and 0.4 mg/ml *E. coli* tRNA. Hybridization proceeded for 16 h at 37 °C and was then washed 5 × with wash solution (50 % formamide, 2x SSC) and 10x with DEPC-treated PBS. Finally, 1 μg/ml DAPI was added to the coverslip, which was then mounted on a glass slide for imaging.

### DNA dye labeling

For live cells, the nucleoid was visualized by incubating 1 μg/ml DAPI with cells in their growth medium for 10 min. Due to a lack of labeling efficiency with DAPI in live cells of some of the species that were studied, all species for Figures 4 and 5 (with the exception of *E. coli* in different conditions) were fixed with 4% formaldehyde for 15 min at room temperature and 30 min on ice. They were then washed three times with 1 × PBS and spun down at 7000 rpm. Fixed *A. excentricus, A. tumefaciens, B. subvibrioides, C. algicola, C. hutchinsonii, H. rosenbergii, B. bacteroides, B. diminuta, C. indologenes, F. johnsoniae, M. xanthus, P. syringae, R. leguminosarum, S. meliloti* and *C. crescentus* were stained with 1 × SYBR Green. Fixed *B. theta, B. ovatus, V. harveyi* and *P. distasonis* were stained with 1 μg/mL DAPI. Fixed *B. megaterium, C. violaceum, J. lividum, V. fischeri, B. subtilis, B. thailandensis, E. coli, Anaerostipes sp., B. xylanisolvens, C. aerofaciens, C. hathewayi, L. reuteri, P. alcalifaciens, R. intestinalis, C. boltae, L. sphaericus* and *P. polymyxa* were stained with 1 μg/ml Hoechst 33342.

### Image processing

Cell outlines were generated using the open-source image analysis software Oufti (Paintdakhi et al., 2016). Nucleoids were detected using Oufti’s objectDetection module. For comparison purposes, we used the same nucleoid detection parameters for all image datasets (see Supplemental Information for parameters), with a single exception (see below). The resulting cell lists were further processed and analyzed in MATLAB (Mathworks) using custom-built algorithms (see below).

One experiment required optimizing our nucleoid detection pipeline. Elongation of *dnaC2* cells under restrictive conditions (related to Figure 2C-D) led to a decrease in the fraction of nucleoid-bound HU-mCherry signal. To overcome this, we used an adjusted nucleoid detection function: Nucleoid_Detection_High_Background.m that uses MATLAB built-in functions to threshold and identify nucleoids within Oufti cell meshes.

SeqA-mCherry signal information was added to *E. coli* cell lists using the MATLAB function Add_SeqA_Area.m (see Supplemental Information for the code). DnaN-CFP information was added to *C. crescentus* cell lists using the MATLAB function Add_DnaN_Area.m.

### Support Vector Machine model for curation of cell contours

In similar fashion as before (Campos et al., 2018), we used an automated approach to identify poor and incorrect cell detections across our datasets. We trained a support vector machine (SVM) model based on 11 normalized phase-contrast features: cell length, cell volume, integrated phase signal, mean cell contour intensity, minimum cell contour intensity, maximum curvature of cell contour, minimum inflated cell contour intensity, mean intensity gradient across the cell contour, maximum variability in contour intensity, mean variability in contour intensity and maximum cell pixel intensity. We visually scored 20,265 cells and used 30% of them (6,080 cells) to train the SVM model. The model was evaluated using a k-fold cross-validation approach, leading to a generalized misclassification rate of 9.9%. We used the remaining 70% of the dataset (14,185 cells) to validate the model. The SVM classifier achieves a balanced classification rate of 90.9% and features an AUROC of 0.9640.

The SVM model underperformed for species (e.g., *C. crescentus*) and mutants (e.g., *dnaC2* at the restrictive temperature) with morphologies that deviated significantly from *E. coli*’s typical rod shape. Therefore, in these instances, we resorted to visual inspection and curation of the obtained cell contours. For time-lapse experiments, visual inspection and manual curation of the cell contours was also required.

### Growth rate measurements

Growth rates were measured in 96-well plates in a Synergy2 microplate reader (BioTek). Cultures were first grown to stationary phase and re-inoculated into 150 μl fresh medium (1/300). Cultures were subsequently grown for 60 h at 37 °C with OD_600_ measurements every 4 min. The maximal growth rate was extracted from the obtained growth curves by fitting the Gompertz function (Zwietering et al., 1990).

### Osmolality measurements

Osmolality of growth media was measured using a Precision Systems 6002 Touch Micro OSMETTE™osmometer, which uses the freezing point method for osmolality measurements. All measurements were conducted in duplicate.

## QUANTIFICATION AND STATISTICAL ANALYSIS

### Cellular characteristics

Properties of individual cells (cell and nucleoid dimensions, DAPI fluorescence intensity, fluorescent marker behavior, etc.) were extracted from cell lists obtained from Oufti using the MATLAB function Extract_Cell_Properties.m. Morphological features (e.g., cell length, width, area, and volume) were determined by summing the dimensions of each individual segment of the cell mesh identified by Oufti. See https://oufti.org/ for more details.

### Correlation coefficients

Kendall correlation coefficients between variables were calculated using MATLAB’s built-in corr function.

### Unconstrained linear fits

Unconstrained linear fits were performed using MATLAB’s built-in polyfit function.

### Nucleoid exclusion of ribosomes

The extent of ribosome exclusion was determined by calculating the signal correlation factor (SCF) between the DNA and ribosome signals. For each individual cell, the SCF was calculated by examining a specific “correlation area”, corresponding to an intracellular region determined by two user-specified parameters. The restriction of the calculation to this area was required to ensure optimal correlation calculations for cells with different shapes and sizes as the smaller cytoplasmic volume at the cell poles and the cell periphery leads to a general decrease in fluorescent signal which, in turn, artificially generates positive biases in the calculation of the SCF. The first parameter was the number of pixels, starting from the cell poles, to exclude from the calculation. The second parameter was the number of pixels, starting from the cell centerline, to include in the calculation. Together, these parameters defined the correlation region for which the correlation between pixel values was determined. Different combinations of these two parameters were scanned for each growth condition and species, and parameters were chosen by finding the minimal average SCF. The minimal average SCF was selected to avoid the positive SCF biases introduced by the cell poles and periphery. The following MATLAB functions were used for this analysis: Pixel_Correlation_Multiple_Experiments_Scan.m, Pixel_Correlation_Parallel.m, Cell_Pixel_Correlation.m, Extract_Cell_Pixels.m, Cell_Projection.m and Taylor_Smooth.m.

### Mean squared displacements of single ribosomal particles

Particle locations determined using the *uTrack* package (Jaqaman et al., 2008) were linked into trajectories based on a previously described algorithm (Crocker and Grier, 1996). Briefly, the most likely trajectories were constructed by minimizing the sum of squared particle displacements between two consecutive frames. Trajectories of lengths smaller than five displacements were removed. Mean squared displacements (MSD) at various time delays were then calculated from individual trajectories. For each frame interval, an ensemble-averaged MSD was obtained by averaging individual MSD curves weighted by the corresponding trajectory lengths. For each MSD curve, the slope was determined by fitting the three smallest time delays using least squares regression and by further dividing by a factor of 4 to obtain the apparent diffusion coefficient D_a_. Similarly, the slope of the log-log MSD curve was determined by fitting the three smallest time delays to obtain the anomalous exponent α. Due to the short average trajectory length, only the three smallest time delays were used to ensure reliable determinations of these values.

### Mean squared displacements of GFP-μNS particles

Cell meshes obtained from Oufti were used to limit particle localization to the region within cells and prevent spurious trajectory linking between cells. Particle localization was performed using the function uNS_Particle_Tracking.m to fit a 2D Gaussian to filtered images.

## DATA AND SOFTWARE AVAILABILITY

All computer code is provided in the Supplemental Information and can also be found at https://github.com/JacobsWagnerLab/published.

