## Supplementary material for "Nucleoid size scaling and intracellular organization of translation across bacteria"

#### Supplemental figure legends

**Figure S1. Related to Figure 1. Cell and nucleoid morphology of *E. coli* cells in different growth media.**

A. Representative phase contrast and DAPI images of *E. coli* cells (CJW6324) grown in liquid cultures of M9 medium supplemented with the indicated carbon source and other chemicals (CAAT: 0.1% casamino acids and 1 µg/ml thiamine) at 37 °C. For a full description of the growth media, see Table S1. Cell contours (green) were generated using Oufti.

B. Bar graph showing the average doubling times of cultures when growing in exponential phase in the indicated growth media. Errors bars indicate the standard deviation between three independent biological replicates. Colors correspond to those used in Figure 1B.

C. Scatter plot of growth medium osmolality versus average NC ratio for *E. coli* cells (CJW6324) grown in the media indicated in B. The color scheme corresponds to the one shown in B. Error bars indicate 95% confidence intervals.

**Figure S2. Related to Figure 1. Variability in DNA content and DAPI staining efficiency does not affect NC ratio measurements in *E. coli*.**

All contour lines represent the 0.10, 0.25, 0.50 and 0.75 probability envelopes of the data.

A. Density contour plots showing the lack of correlation between the NC ratio and the normalized DAPI signal intensity (total intensity divided by cell area) for CJW6324 cells grown in the indicated growth media (for a full description of the growth media, see Table S1).

B. Density contour plots showing the lack of correlation between total DAPI signal intensity and NC ratio for CJW6324 cells grown in the indicated growth media.

**Figure S3. Related to Figure 1. The nucleoid-labeling method does not affect NC ratio measurements and nucleoid scaling occurs independently of cell division.**

A. Density contour plots showing the strong correlation between cell area and nucleoid area for the different strains and nucleoid-labeling methods. The contour lines represent the 0.10, 0.25, 0.50 and 0.75 probability envelopes of the data. *E. coli* cells expressing fluorescent fusions to the α or β HU-subunit (CJW5556 producing HupA-mCherry and CJW4656 producing HupB-CFP) were grown in M9 medium supplemented with 0.2% glycerol, 0.1% casamino acids, and 1 μg/ml thiamine, and either stained with 1 mM DAPI or not. For DAPI-stained cultures, nucleoid areas were determined using both the DAPI signal (first column) and the signal from the fluorescently-labeled HU (second column). For cultures not stained with DAPI, only the HU-CFP or HU-mCherry fluorescence signal was used (third column). As a reference, the relationship between cell area and nucleoid area for wild-type *E. coli* MG1655 cells grown in the same conditions, and stained with DAPI, is shown in blue (fourth column).

B. Average NC ratio (with error bars representing the standard deviation) for experiments shown in A.

C. Scatter plot of nucleoid area (measured using DAPI signal) versus nucleoid area (measured using fluorescently labeled HU) for individual cells. Dotted red line represents the identity line, which indicates a perfect agreement between the two signals.

D. Plot showing the average nucleoid area per cell area bin for untreated and cephalexin-treated *E. coli* cells (CJW4677) grown in the indicated growth media (for a full description of the growth media, see Table S1) and stained with DAPI. Error bars represent the standard deviation of the nucleoid area for each cell area bin.

**Figure S4. Related to Figure 2. Nucleoid size scaling occurs independently of DNA replication.**

A. Density scatter plot of relative cell age versus total DAPI signal of *E. coli* cells (CJW6324) grown in M9 medium supplemented with 0.2% glycerol. Relative cell age was calculated by ordering cells according to their cell area and using the relationship described by Wold et al. (1994). Using this methodology, the overlap in cell size between cells in the B, C and D cell cycle periods (see Figure 2B) leads to an almost monotonous increase in total DAPI intensity with cell age. This precludes the use of this method for identifying cells in different cell cycle periods. The gray scale indicates the relative density of dots (cells) in a given area of the chart. Red dots indicate the mean total DAPI intensity values for cell age bins whereas the error bars show the standard deviations.

B. Bar graph showing the average NC ratio of CJW6324 cells in the B, C, or D cell cycle periods. The cells were grown in the indicated growth media (for a full description of the growth media, see Table S1).

C. Bar graph showing the slope of an unconstrained linear fit between cell area and nucleoid area for the cells shown in panel B.

D. Density contour plots of cell area versus nucleoid area for CJW6324 cells in B, C, and D periods. Growth media are indicated on each panel. The contour lines represent the 0.10, 0.25, 0.50 and 0.75 probability envelopes of the data. The nucleoid was detected by DAPI staining.

**Figure S5. Related to Figure 3. Robust nucleoid size scaling in *C. crescentus*.**

All contour lines represent the 0.10, 0.25, 0.50 and 0.75 probability envelopes of the data.

A. Density contour plots showing the lack of correlation between the NC ratio and the total DAPI intensity (left) and between the NC ratio and the normalized DAPI intensity (right) for *C. crescentus* CB15N cells grown in M2 medium supplemented with 0.2% glucose (M2G).

B. Density contour plots of cell area versus nucleoid area for CJW5806 cells expressing an mCherry fusion to the HU subunit HU2. Cells were grown in M2G, and the nucleoid area was determined using the HU-mCherry fluorescence signal.

C. Density contour plots of cell area versus nucleoid area for CB15N cells grown in M2G or PYE. For a full description of the growth media, see Table S1. The nucleoid was detected by DAPI staining.

D. Density contour plots showing the strong correlation between cell area and nucleoid area for the indicated *C. crescentus* strains. All cells, whether FtsZ-depleted (CJW3821) or carrying a Δ*rodZ* (CJW1842) and Δ*hfq* (CJW5477) deletion, were grown in PYE, and the nucleoid was detected by DAPI staining. For FtsZ depletion, CJW3821 cells grown in PYE that contained xylose (0.3%) were washed and resuspended into PYE lacking xylose to inhibit *ftsZ* expression. Cells were imaged 3-6 h after cell resuspension. The nucleoid was detected by DAPI staining.

E. Average NC ratio (with error bars representing the standard deviation) for all strains and experiments included in this figure. The nucleoid was detected by DAPI staining.

F. Density contour plots showing the strong correlation between cell area and nucleoid area for cells carrying a *popZ* deletion (CJW2238). The cells were grown in M2G because growth in PYE results in severe cell filamentation and loss of viability (Bowman et al., 2008; Ebersbach et al., 2008). The nucleoid was detected by DAPI staining.

**Figure S6. Related to Figure 4. Cell and nucleoid morphology across bacterial species.**
A. Representative phase contrast and DAPI images of the indicated species grown in the media detailed in the STAR methods section. The images were processed using Oufti to identify the contours of the formaldehyde-fixed cells (green) and stained nucleoids (purple).
B. Scatter plot showing the lack of correlation between genome size and the average NC ratio (left), the average nucleoid area (middle) or the average cell volume (right) for the different species.

**Figure S7. Related to Figure 4. Nucleoid-staining efficiency does not affect NC ratio measurements.**
All contour lines represent the 0.10, 0.25, 0.50 and 0.75 probability envelopes of the data.
A. Density contour plots showing the lack of correlation between normalized DNA signal intensity (total signal intensity divided by the cell area) and NC ratio for the indicated species.
B. Density contour plots showing the lack of correlation between total DNA signal intensity and NC ratio for the indicated species.
C. Scatter plot showing the lack of correlation between the average NC ratio and the average normalized DNA signal intensity for all included species. Abbreviated species names are indicated next to the corresponding datapoint; see Key Resources Table for a full name description.

**Figure S8. Related to Figures 6 and 7. GFP-μNS particle trajectories, log-log MSD plots and calculation of the signal correlation factor in live and fixed cells.**
A. Randomly picked trajectories of GFP-μNS particles belonging to each of the intensity bins indicated in Figure 6A. *E. coli* cells (CJW6723) were grown in M9 medium supplemented with 0.2% glycerol and *C. crescentus* cells (CJW6917) in M2 medium supplemented with 0.2% glucose.
B. Same plot as Figure 6B on a log-log scale.
C. Same plots as Figure 6C-D on a log-log scale.
D. Schematic showing the area used for calculating the signal correlation factor (SCF). Two parameters define the "correlation area" to ensure optimal SCF calculations for cells with different shapes and sizes. The first parameter is the number of pixels, starting from the cell poles, to exclude from the calculation. The second parameter is the number of pixels, starting from the cell centerline, to include in the calculation. The SCF is calculated as the Pearson correlation between the pixel values of the two signals within this correlation area.
E. Left, frequency distribution of SCF values between nucleoid and ribosome signals in *E. coli* cells (CJW4677) after treatment with the indicated concentrations of formaldehyde for 15 min at room

temperature followed by 30 min on ice for fixation. The nucleoid was detected by DAPI staining, and the ribosome via the GFP fusion to the ribosomal protein RlpA. Cells were grown in M9 medium supplemented with 0.2% glycerol, 0.1% casamino acids, and 1μg/ml thiamine. Right, average SCF values for nucleoid and ribosome signals for the populations of cells shown in B.

#### Supplemental movie legends

**Movie S1. Related to Figure 6. GFP-μNS particle mobility in *E. coli*.**

The movie shows a time-lapse sequence of a GFP-μNS particle moving in an *E. coli* cell (CJW6723) grown in M9 medium supplemented with 0.2% glycerol at 30 °C. The time-lapse sequence is shown as an overlay of fluorescence images (green) with the corresponding phase contrast images. The spot intensity (hence size) of the GFP-μNS particle is comparable to that of the GFP-μNS particle in the *C. crescentus* cell in Movie S2.

**Movie S2. Related to Figure 6. GFP-μNS particle mobility in *C. crescentus*.**

The movie shows a time-lapse sequence of a GFP-μNS particle moving in a *C. crescentus* cell (CJW6917) grown in M2 medium supplemented with 0.2% glucose at 30 °C. The time-lapse sequence is shown as an overlay of fluorescence images (green) with the corresponding phase contrast images. The spot intensity (hence size) of the GFP-μNS particle is comparable to that of the GFP-μNS particle in the *E. coli* cell in Movie S1.

**Movie S3. Related to Figure 6. Single-molecule detection of fluorescently labeled ribosomes in *E. coli*.**

The movie shows a representative time-lapse sequence of the detection of individual ribosomes (labeled with RpsV-mEos2) in *E. coli* (SX289). The measurement frame interval is 50 ms.

**Movie S4. Related to Figure 6. Single-molecule detection of fluorescently labeled ribosomes in *C. crescentus*.**

The movie shows a representative time-lapse sequence of the detection of individual ribosomes (labeled with RplA-Dendra2) in *C. crescentus* (CJW5156). The measurement frame interval is 50 ms.

**Movie S5. Related to Figure 6. FRAP microscopy in *E. coli* producing fluorescently labeled ribosomes.**

The movie shows a representative time-lapse sequence of the evolution of the ribosomal fluorescence recovery over time (up to 450 s) along the length of a cephalexin-treated *E. coli* cell (CJW4677) following photobleaching of about half of the cell. Same cell as in Figure 6G.

**Movie S6. Related to Figure 6. FRAP microscopy in *C. crescentus* producing fluorescently labeled ribosomes.**
The movie shows a representative time-lapse sequence of the evolution of the ribosomal fluorescence recovery over time (up to 450 s) along the length of an FtsZ-depleted cephalexin-treated *C. crescentus* cell (CJW3821) following photobleaching of about half of the cell. Same cell as in Figure 6G.

### Supplementary tables

**Table S1. Related to Figure 1, Figure 8 and STAR methods. Growth medium composition.**

| Abbreviation | Medium composition |
|---|---|
| 1 x M9 salts | 6 g/l $Na_2HPO_4 \cdot 7H_2O$, 3 g/l $KH_2PO_4$, 0.5 g/l NaCl, 1 g $NH_4Cl$, 2 mM $MgSO_4$, 0.1 mM $CaCl_2$ |
| M9acet | 1x M9 salts + 0.2% acetate (sodium salt) |
| M9αKG | 1x M9 salts + 0.2% α-ketoglutarate (sodium salt) |
| M9fum | 1x M9 salts + 0.2% fumarate (disodium salt) |
| M9glu | 1x M9 salts + 0.2% glucose |
| M9gly | 1x M9 salts + 0.2% glycerol |
| M9lact | 1x M9 salts + 0.2% lactose |
| M9Lara | 1x M9 salts + 0.2% L-arabinose |
| M9mala | 1x M9 salts + 0.2% malate (sodium salt) |
| M9malt | 1x M9 salts + 0.2% maltose |
| M9mann | 1x M9 salts + 0.2% mannose |
| M9pyr | 1x M9 salts + 0.2% pyruvate (sodium salt) |
| M9sorb | 1x M9 salts + 0.2% sorbitol |
| M9succ | 1x M9 salts + 0.2% succinate (disodium salt) |
| M9treh | 1x M9 salts + 0.2% trehalose |
| M9xyl | 1x M9 salts + 0.2% xylose |
| M9acetCAAT | 1x M9 salts + 0.2% acetate + 0.1% casamino acids + 1 µg/ml thiamine |

| | |
|---|---|
| M9αKGCAAT | 1x M9 salts + 0.2% α-ketoglutarate + 0.1% casamino acids + 1 μg/ml thiamine |
| M9fumCAAT | 1x M9 salts + 0.2% fumarate + 0.1% casamino acids + 1 μg/ml thiamine |
| M9gluCAAT | 1x M9 salts + 0.2% glucose + 0.1% casamino acids + 1 μg/ml thiamine |
| M9glyCAAT | 1x M9 salts + 0.2% glycerol + 0.1% casamino acids + 1 μg/ml thiamine |
| M9lactCAAT | 1x M9 salts + 0.2% lactose + 0.1% casamino acids + 1 μg/ml thiamine |
| M9LaraCAAT | 1x M9 salts + 0.2% L-arabinose + 0.1% casamino acids + 1 μg/ml thiamine |
| M9malaCAAT | 1x M9 salts + 0.2% malate + 0.1% casamino acids + 1 μg/ml thiamine |
| M9maltCAAT | 1x M9 salts + 0.2% maltose + 0.1% casamino acids + 1 μg/ml thiamine |
| M9mannCAAT | 1x M9 salts + 0.2% mannose + 0.1% casamino acids + 1 μg/ml thiamine |
| M9pyrCAAT | 1x M9 salts + 0.2% pyruvate + 0.1% casamino acids + 1 μg/ml thiamine |
| M9sorbCAAT | 1x M9 salts + 0.2% sorbitol + 0.1% casamino acids + 1 μg/ml thiamine |
| M9succCAAT | 1x M9 salts + 0.2% succinate + 0.1% casamino acids + 1 μg/ml thiamine |
| M9trehCAAT | 1x M9 salts + 0.2% trehalose + 0.1% casamino acids + 1 μg/ml thiamine |
| M9xylCAAT | 1x M9 salts + 0.2% xylose + 0.1% casamino acids + 1 μg/ml thiamine |
| LB | 10 g/l NaCl, 5 g/l yeast extract, 10 g/l tryptone |
| LBS | 20 g/l NaCl, 5 g/l yeast extract, 10 g/l tryptone |
| PYE | 2 g/l bacto-peptone, 1 g/l yeast extract, 1 mM $MgSO_4$, 0.5 mM $CaCl_2$ |
| M2G | 0.87 g/l $Na_2HPO_4$, 0.54 g/l $KH_2PO_4$, 0.50 g/l $NH_4Cl$, 0.2% glucose, 0.5 mM $MgSO_4$, 0.5 mM $CaCl_2$, 0.01 mM $FeSO_4$ |
| CYE | 1% casitone, 0.5% yeast extract, 10 mM MOPS (pH 7.6) and 4 mM MgSO4 |
| DSMZ Medium 172 | 1 g/l yeast extract, 1g/l tryptone, 24.7 g/l NaCl, 0.7 g/l KCl, 6.3 g/l $MgSO_4 \cdot 7H_2O$, 4.6 g/l $MgCl_2 \cdot 6H_2O$, 1.2 g/l $CaCl_2 \cdot 2H_2O$, 0.2 g/l $NaHCO_3$ |
| Nutrient broth medium | 3 g/l beef extract, 5 g/l peptone |
| TYG | 10 g/l tryptone peptone, 5 g/l bacto yeast extract, 2 g/l D-glucose, 500 mg L-cysteine, 1 ml vitamin K solution (1 mg/ml stock solution), 1 ml/l 0.8% $CaCl_2$, 1 ml $FeSO_4$ solution (0.4 mg/ml stock solution), 1 ml/l histidine hematin solution (200 mM stock solution), 4 mL resazurin solution (0.25 g/ml stock solution), 100mM potassium phosphate buffer (pH 7.2), 20 mg/l $MgSO_4$-$7H_2O$, 400 mg/l $NaHCO_3$, 80 mg/l NaCl |
| GMM | 2 g/l tryptone peptone, 1 g/l yeast extract, 0.4 g/l D-glucose, 0.5 g/l L-cysteine, 1 g/l cellobiose, 1 g/l maltose, 1 g/l fructose, 5 g/l meat extract, 100 mM |

| | |
|---|---|
| | $KH_2PO_4$, 0.008 mM $MgSO_4$-$7H_2O$, 4.8 mM $NaHCO_3$, 1.37 mM NaCl, 8 mg/l $CaCl_2$, 5.8 mM menadione (vitamin K), 1.44 mM $FeSO_4$, 1 ml/l histidine hematin solution (200 mM stock solution), 2 ml Tween 80 (25% stock solution), 10 ml ATCC vitamin mix, 10 mL ATCC trace mineral mix, 30 mM acetic acid, 1 mM isovaleric acid, 8 mM propionic acid, 4 mM butyric acid, 4 mM resazurin |

**Table S2. Related to STAR methods. Construction of strains and plasmids used in this study.**

All constructed strains and plasmids were verified by both PCR and sequencing.

| Strain or plasmid | Construction method |
|---|---|
| Strains | |
| CJW6324 | This strain was constructed using *E. coli* K12 MG1655 as a parental strain. An *ftsZ-venus*$^{SW}$*-lpxC*-FRT-*kan*-FRT-*secM'* amplicon was generated by PCR using oligonucleotides SG25 and SG60, and pKD13-ftsZ-venus$^{SW}$-lpxC-FRT-kan-FRT-secM' as a template. This amplicon was used to replace the native *ftsZ* gene of MG1655 by lambda Red-mediated recombination (Datsenko and Wanner, 2000), yielding the chromosomal *ftsZ-venus*$^{SW}$ fusion. The kanamycin resistance cassette was subsequently excised by transiently equipping this strain with plasmid pCP20 expressing the Flp site-specific recombinase (Cherepanov and Wackernagel, 1995). Subsequently, an *mcherry*-FRT-*cat*-FRT amplicon flanked by short (50 bp) nucleotide sequences homologous to its target region (50 nt on both sides of the *seqA* stop codon) was generated by PCR amplification using oligonucleotides SG5 and SG6, and pKD3-mCherry as a template. This amplicon was used to replace the stop codon of the native *seqA* gene, yielding the chromosomal *seqA-mcherry* fusion. The chloramphenicol resistance cassette was subsequently excised by transiently equipping this strain with plasmid pCP20 expressing the Flp site-specific recombinase (Cherepanov and Wackernagel, 1995). |
| CJW4656 | The *hupB-cfp-kan* fragment from *E. coli* CSH50 *hupB*::*hupB-cfp-kan* (Berger et al., 2010) was moved into *E. coli* K12 MG1655 by P1 transduction. |
| CJW4677 | Strain CJW4677 was constructed using *E. coli* K12 MG1655 as a parental strain. A pGFPC2*gfp*-FRT-*cm*-FRT plasmid was generated using oligonucleotides WTG1 and WTG2 to amplify the FRT-*cm*-FRT region from pKD3 (Datsenko and Wanner, 2000) followed by insertion into the NheI site of pGFPC2. An *rplA*-*gfp*-FRT-*cm*-FRT |

| | |
|---|---|
| | amplicon was created using oligonucleotides WTG3 and WTG4. This amplicon was used to replace the native *rplA* gene of MG1655 by lambda Red-mediated recombination (Datsenko and Wanner, 2000), yielding the chromosomal *rplA-gfp* fusion. The kanamycin resistance cassette was subsequently excised by transiently equipping this strain with plasmid pCP20 expressing the Flp site-specific recombinase (Cherepanov and Wackernagel, 1995). |
| CJW6370 | The *hupA-mcherry* fusion (Marceau et al., 2011) was moved into *E. coli* K12 MG1655 *dnac2* (Lobner-Olesen et al., 2008) by P1 transduction. |
| CJW6723 | The kanamycin resistance cassette was excised from CJW4617 (Parry et al., 2014) by transiently equipping this strain with plasmid pCP20 expressing the Flp site-specific recombinase (Cherepanov and Wackernagel, 1995). The *hupA-mcherry* fusion (Marceau et al., 2011) was subsequently moved into this strain by P1 transduction. |
| CJW6917 | Genomic DNA from CJW4617 (Parry et al., 2014) was amplified with oligonucleotides WTG5 and WTG6, which was then inserted into the KpnI/NheI sites in pXGFPN-2 (pMT582) (Thanbichler et al., 2007). The resulting plasmid was transformed into CB15N for integration into the xylose operon to allow inducible expression of GFP-μNS. |
| Plasmids | |
| pKD13-ftsZ-venus$^{SW}$-lpxC-FRT-kan-FRT-secM' | The FtsZ-Venus sandwich fusion (Moore et al., 2017) was obtained from Harold Erickson (Duke University). The entire fusion and its neighboring *lpxC* gene (including its terminator) were PCR amplified (using oligonucleotides SG28 and SG41) and integrated, by Gibson assembly (Gibson et al., 2009), into plasmid pKD13 (Datsenko and Wanner, 2000), which was linearized by PCR amplification using oligonucleotides SG43 and SG36 (each containing an overhang of 30 nt, overlapping with the ends with the *ftsZ-Venus$^{SW}$-lpxC* amplicon). The obtained plasmid was subsequently linearized by PCR amplification using oligonucleotides SG61 and SG62 (each containing an overhang of 30 nt, overlapping with the ends of the amplicon to be inserted).An amplicon containing a truncated version of *secM* (*secM'*) was obtained by PCR amplification using oligonucleotides SG59 and SG60 and integrated, by Gibson assembly (Gibson et al., 2009), into the linearized plasmid, yielding pKD13-ftsZ-venus$^{SW}$-lpxC-FRT-kan-FRT-secM'. |

| | |
|---|---|
| pKD3-mCherry | An *mcherry* amplicon was obtained from pTU136 (Uehara et al., 2009) using oligonucleotides SG1 and SG2. In addition to amplifying the *mcherry* sequence, this primer pair also added a flexible linker (encoding Gly-Ser-Gly-Ser-Gly-Ser), facilitating folding of fluorescent fusion proteins constructed with this fluorescent protein. The entire amplicon was subsequently integrated, by Gibson assembly (Gibson et al., 2009), into plasmid pKD3 (Datsenko and Wanner, 2000), linearized by PCR amplification using oligonucleotides SG3 and SG4 (each containing an overhang of 30 nt, overlapping with the ends with the *mcherry* amplicon), yielding pKD3-mCherry. |

**Table S3. Related to STAR methods. Oligonucleotides used in this study.**

| Name | Sequence (5’ to 3’) |
|---|---|
| SG28 | ATGTTTGAACCAATGGAACTTACC |
| SG41 | TTAAGAAAACAGCGTTCGCACC |
| SG36 | GTCATTGGTAAGTTCCATTGGTTCAAACATAATCGCTCAAGACGTGTAATGCTG |
| SG43 | TAAAAAACGGTGCGAACGCTGTTTTCTTAAGTGTAGGCTGGAGCTGCTTCGA |
| SG59 | GCACTTTTCCGCACAACTTATC |
| SG60 | ACCCTTTTCAGACGGCGTG |
| SG61 | ACCCAGGAAGGCACGCCGTCTGAAAAGGGTTAATTCTCATGTTTGACAGCTTATCACTG |
| SG62 | CGAATGAAGATAAGTTGTGCGGAAAAGTGCATTCCGGGGATCCGTCGACC |
| SG25 | CCATAAACTGCCAGGCATCAA |
| WTG1 | ACGTGCTAGCGTGTAGGCTGGAGCTGCTTC |
| WTG2 | ACGTGCTAGCATGGGAATTAGCCATGGTCC |
| WTG3 | CCATGGGTGCAGGTGTTGCAGTTGACCAGGCTGGCCTGAGCGCTTCTGTAAACCGGTCGGCCACCATGGTGAGC |
| WTG4 | GCATTATACGTGGGGGTAAGATTGTAGACAAAATCACCGCCCACGTAAAGGCTCCTGCAGCCCGGGGGATCC |
| WTG5 | TAATTAATATGCATGGTACCTCCAGTGACATGGTAGACGGGATTA |
| WTG6 | TCCCCCGGGCTGCAGCTAGCTCAGGCTGAAAATCTTCTCTCATCC |

### Supplemental information references


Baba, T., Ara, T., Hasegawa, M., Takai, Y., Okumura, Y., Baba, M., Datsenko, K.A., Tomita, M., Wanner, B.L., and Mori, H. (2006). Construction of *Escherichia coli* K-12 in-frame, single-gene knockout mutants: the Keio collection. Mol Syst Biol *2*, 2006 0008.

Berger, M., Farcas, A., Geertz, M., Zhelyazkova, P., Brix, K., Travers, A., and Muskhelishvili, G. (2010). Coordination of genomic structure and transcription by the main bacterial nucleoid-associated protein HU. EMBO Rep *11*, 59-64.

Bowman, G.R., Comolli, L.R., Zhu, J., Eckart, M., Koenig, M., Downing, K.H., Moerner, W.E., Earnest, T., and Shapiro, L. (2008). A polymeric protein anchors the chromosomal origin/ParB complex at a bacterial cell pole. Cell *134*, 945-955.

Cherepanov, P.P., and Wackernagel, W. (1995). Gene disruption in Escherichia coli: TcR and KmR cassettes with the option of Flp-catalyzed excision of the antibiotic-resistance determinant. Gene *158*, 9-14.

Datsenko, K.A., and Wanner, B.L. (2000). One-step inactivation of chromosomal genes in *Escherichia coli* K-12 using PCR products. Proc Natl Acad Sci U S A *97*, 6640-6645.

Ebersbach, G., Briegel, A., Jensen, G.J., and Jacobs-Wagner, C. (2008). A self-associating protein critical for chromosome attachment, division, and polar organization in *Caulobacter*. Cell *134*, 956-968.

Gibson, D.G., Young, L., Chuang, R.Y., Venter, J.C., Hutchison, C.A., 3rd, and Smith, H.O. (2009). Enzymatic assembly of DNA molecules up to several hundred kilobases. Nat Methods *6*, 343-345.

Lobner-Olesen, A., Slominska-Wojewodzka, M., Hansen, F.G., and Marinus, M.G. (2008). DnaC inactivation in *Escherichia coli* K-12 induces the SOS response and expression of nucleotide biosynthesis genes. PLoS One *3*, e2984.

Marceau, A.H., Bahng, S., Massoni, S.C., George, N.P., Sandler, S.J., Marians, K.J., and Keck, J.L. (2011). Structure of the SSB-DNA polymerase III interface and its role in DNA replication. EMBO J *30*, 4236-4247.

Moore, D.A., Whatley, Z.N., Joshi, C.P., Osawa, M., and Erickson, H.P. (2017). Probing for Binding Regions of the FtsZ protein surface through site-directed insertions: discovery of fully functional FtsZ-fluorescent proteins. J Bacteriol *199*.

Parry, B.R., Surovtsev, I.V., Cabeen, M.T., O'Hern, C.S., Dufresne, E.R., and Jacobs-Wagner, C. (2014). The bacterial cytoplasm has glass-like properties and is fluidized by metabolic activity. Cell *156*, 183-194.

Thanbichler, M., Iniesta, A.A., and Shapiro, L. (2007). A comprehensive set of plasmids for vanillate- and xylose-inducible gene expression in *Caulobacter crescentus*. Nucleic Acids Res *35*, e137.

Uehara, T., Dinh, T., and Bernhardt, T.G. (2009). LytM-domain factors are required for daughter cell separation and rapid ampicillin-induced lysis in *Escherichia coli*. J Bacteriol *191*, 5094-5107.

Wold, S., Skarstad, K., Steen, H.B., Stokke, T., and Boye, E. (1994). The initiation mass for DNA replication in *Escherichia coli* K-12 is dependent on growth rate. EMBO J *13*, 2097-2102.

### KEY RESOURCES TABLE

| REAGENT or RESOURCE | SOURCE | IDENTIFIER |
|---|---|---|
| Bacterial and Virus Strains | | |
| *E. coli* MG1655 | (Guyer et al., 1981) | (Ec) |
| *E. coli* MG1655 *hupB*::*hupB-cfp* | This work | CJW4656 |
| *E. coli* MG1655 *rplA*::*rplA-gfp* | This work | CJW4677 |
| *E. coli* BW25113 *hupA*::*hupA-mcherry* | (Paintdakhi et al., 2016) | CJW5556 |
| *E. coli* MG1655 *seqA*::*seqA-mcherry ftsZ*::*ftsZ-venus*$^{SW}$ | This work | CJW6324 |
| *E. coli* MG1655 *dnaC2 hupA*::*hupA-mcherry* | This work | CJW6370 |
| *E. coli* MG1655 *PlacOAYZ*::*Plac-gfp-μNS hupA*::*hupA-mcherry* | This work | CJW6723 |
| *E. coli* BW25993 *rpsB*::*rpsB-meos2* | (Sanamrad et al., 2014) | CJW6769 |
| *E. coli* BW25993 *rpsV*::*rpsV-meos2* | (Wang et al., 2011) | SX289 |
| *C. crescentus* CB15N | (Evinger and Agabian, 1977) | (Cc) |
| *C. crescentus* CB15N *rodZ*::Himar*1* | (Alyahya et al., 2009) | CJW1842 |
| *C. crescentus* CB15N *popZ*::omega | (Ebersbach et al., 2008) | CJW2238 |
| *C. crescentus* CB15N *ftsZ*::Pxyl-ftsZ *rplA*::pL1-GFPC-1 | (Montero Llopis et al., 2012) | CJW3821 |
| *C. crescentus* CB15N *rplA*::pL1-dendra2 | (Lim et al., 2014) | CJW5156 |
| *C. crescentus* CB15N *hfq*::*tet* | (Irnov et al., 2017) | CJW5477 |
| *C. crescentus* CB15N *hu2*::pCHYC2-hu2' | (Arias-Cartin et al., 2017) | CJW5806 |
| *C. crescentus* CB15N *hu2*::pCHYC2-hu2' dnaN::pCFPC1-dnaNend' | (Arias-Cartin et al., 2017) | CJW5969 |
| *C. crescentus* CB15N *xylX*::pXyl-egfp-μNS-kan | This work | CJW6917 |
| *Rhizobium leguminosarum* bv. *trifolii* R200 | Nora Ausmees | CJW355 (Rl) |
| *Sinorhizobium meliloti* 1021 | Jacques Batut | CJW356 (Sm) |
| *Pseudomonas syringae* pv. syringae B728a | Steven Lindow | CJW410 (Ps) |
| *Agrobacterium tumefaciens* 3101 (*Rhizobium rodiobacter*) | Savithramma Dinesh-Kumar | CJW501 (At) |
| *Asticcacaulis excentricus* ATCC 15261 | Jeanne S. Poindexter | CJW960 (Ae) |
| *Chryseobacterium indologenes* | Jo Handelsman | CJW4422 (Ci) |
| *Janthinobacterium lividum* | Jo Handelsman | CJW4423 (Jl) |
| *Vibrio harveyi* BB120 | Bonnie Bassler | CJW5482 (Vh) |
| *Burkholderia thailandensis* E264 | Peter Greenberg | CJW5484 (But) |
| *Myxococcus xanthus* DZF1 | David Zusman | CJW5485 (Mx) |
| *Bacillus subtilis* subsp. *subtilis* str. NCIB 3610 | Wade Winkler | CJW5495 (Bs) |
| *Brevundimonas subvibrioides* | Pamela Brown | CJW5565 (Brs) |
| *Brevundimonas bacteroides* | Pamela Brown | CJW5566 (Brb) |
| *Brevundimonas diminuta* | Pamela Brown | CJW5571 (Brd) |

| | | |
|---|---|---|
| *Hirschia rosenbergii* | Pamela Brown | CJW5574 (Hr) |
| *Vibrio fischeri* ES114 | Eric Stabb | CJW5630 (Vf) |
| *Paenibacillus polymyxa* ATCC 7070 | ATCC Bacteriology Collection | CJW5750 (Pp) |
| *Lysinibacillus sphaericus* ATCC 4525 | ATCC Bacteriology Collection | CJW5751 (Ls) |
| *Bacillus megaterium* ATCC 14581 | ATCC Bacteriology Collection | CJW5752 (Bm) |
| *Flavobacterium johnsoniae* ATCC 17061$^T$ | Mark McBride | CJW5841 (Fj) |
| *Cytophaga hutchinsonii* ATCC 33406 (glucose-adapted) | Mark McBride (Zhu and McBride, 2014) | CJW5842 (Cyh) |
| *Cellulophaga algicola* IC166$^T$ | Mark McBride | DSM14237 (Ca) |
| *Chromobacterium violaceum* | Jo Handelsman | N/A (Cv) |
| *Bacteroides ovatus* | Andrew Goodman | N/A (Bt) |
| *Bacteroides thetaiotaomicron* | Andrew Goodman | N/A (Bo) |
| *Bacteroides xylanisolvens* | Andrew Goodman | N/A (Bx) |
| *Parabacteroides distasonis* | Andrew Goodman | N/A (Pd) |
| *Providencia alcalifaciens* | Andrew Goodman | N/A (Pa) |
| *Roseburia intestinalis* | Andrew Goodman | N/A (Ri) |
| *Anaerostipes sp.* | Andrew Goodman | N/A (As) |
| *Clostridium boltae* | Andrew Goodman | N/A (Cb) |
| *Clostridium hathewayi* | Andrew Goodman | N/A (Ch) |
| *Lactoacillus reuteri* | Andrew Goodman | N/A (Lr) |
| *Collinsella aerofaciens* | Andrew Goodman | N/A (Coa) |
| Chemicals, Peptides, and Recombinant Proteins | | |
| 4',6-Diamidine-2'-phenylindole dihydrochloride (DAPI) fluorescent dye | Thermo Fisher Scientific | Cat#D1306 |
| Hoechst 33342 fluorescent dye | Thermo Fisher Scientific | Cat#H3570 |
| SYBR Green fluorescent dye | Thermo Fisher Scientific | Cat#S7564 |
| Agarose | AmericBio | Cat#AB00972-00500 |
| T5 exonuclease | New England Biolabs | Cat#M0363S |
| Taq DNA ligase | New England Biolabs | Cat#M0208L |
| Phusion high-fidelity polymerase | New England Biolabs | Cat#M0530S |
| DpnI restriction enzyme | New England Biolabs | Cat#R0176S |
| NheI restriction enzyme | New England Biolabs | Cat#R0131S |
| KpnI restriction enzyme | New England Biolabs | Cat#R0142S |
| Cephalexin hydrate | Sigma Aldrich | Cat#C4895 |
| LB medium | Fisher Scientific | Cat#BP1426-2 |
| Oligonucleotides | | |
| Primers for strain construction | This work, Integrated DNA Technologies | See Table S3 |
| Recombinant DNA | | |
| pKD13-ftsZ-venus$^{SW}$-lpxC-FRT-kan-FRT-secM' | This work | N/A |
| pKD3-mCherry | This work | N/A |
| Software and Algorithms | | |

| | | |
|---|---|---|
| MATLAB | Mathworks | |
| Oufti | (Paintdakhi et al., 2016) | https://oufti.org/ |
| ImageJ | (Collins, 2007) | https://imagej.nih.gov/ij/ |

#### Key Resources Table References


Alyahya, S.A., Alexander, R., Costa, T., Henriques, A.O., Emonet, T., and Jacobs-Wagner, C. (2009). RodZ, a component of the bacterial core morphogenic apparatus. Proc Natl Acad Sci U S A *106*, 1239-1244.

Arias-Cartin, R., Dobihal, G.S., Campos, M., Surovtsev, I.V., Parry, B., and Jacobs-Wagner, C. (2017). Replication fork passage drives asymmetric dynamics of a critical nucleoid-associated protein in *Caulobacter*. EMBO J *36*, 301-318.

Collins, T.J. (2007). ImageJ for microscopy. Biotechniques *43*, 25-30.

Ebersbach, G., Briegel, A., Jensen, G.J., and Jacobs-Wagner, C. (2008). A self-associating protein critical for chromosome attachment, division, and polar organization in *Caulobacter*. Cell *134*, 956-968.

Evinger, M., and Agabian, N. (1977). Envelope-associated nucleoid from *Caulobacter crescentus* stalked and swarmer cells. J Bacteriol *132*, 294-301.

Guyer, M.S., Reed, R.R., Steitz, J.A., and Low, K.B. (1981). Identification of a sex-factor-affinity site in *E. coli* as gamma delta. Cold Spring Harb Symp Quant Biol *45 Pt 1*, 135-140.

Irnov, I., Wang, Z., Jannetty, N.D., Bustamante, J.A., Rhee, K.Y., and Jacobs-Wagner, C. (2017). Crosstalk between the tricarboxylic acid cycle and peptidoglycan synthesis in *Caulobacter crescentus* through the homeostatic control of alpha-ketoglutarate. PLoS Genet *13*, e1006978.

Lim, H.C., Surovtsev, I.V., Beltran, B.G., Huang, F., Bewersdorf, J., and Jacobs-Wagner, C. (2014). Evidence for a DNA-relay mechanism in ParABS-mediated chromosome segregation. Elife *3*, e02758.

Montero Llopis, P., Sliusarenko, O., Heinritz, J., and Jacobs-Wagner, C. (2012). *In vivo* biochemistry in bacterial cells using FRAP: insight into the translation cycle. Biophys J *103*, 1848-1859.

Paintdakhi, A., Parry, B., Campos, M., Irnov, I., Elf, J., Surovtsev, I., and Jacobs-Wagner, C. (2016). Oufti: an integrated software package for high-accuracy, high-throughput quantitative microscopy analysis. Mol Microbiol *99*, 767-777.

Sanamrad, A., Persson, F., Lundius, E.G., Fange, D., Gynna, A.H., and Elf, J. (2014). Single-particle tracking reveals that free ribosomal subunits are not excluded from the *Escherichia coli* nucleoid. Proc Natl Acad Sci U S A *111*, 11413-11418.

Wang, W., Li, G.W., Chen, C., Xie, X.S., and Zhuang, X. (2011). Chromosome organization by a nucleoid-associated protein in live bacteria. Science *333*, 1445-1449.

Zhu, Y., and McBride, M.J. (2014). Deletion of the *Cytophaga hutchinsonii* type IX secretion system gene *sprP* results in defects in gliding motility and cellulose utilization. Appl Environ Microbiol *98*, 763-775.

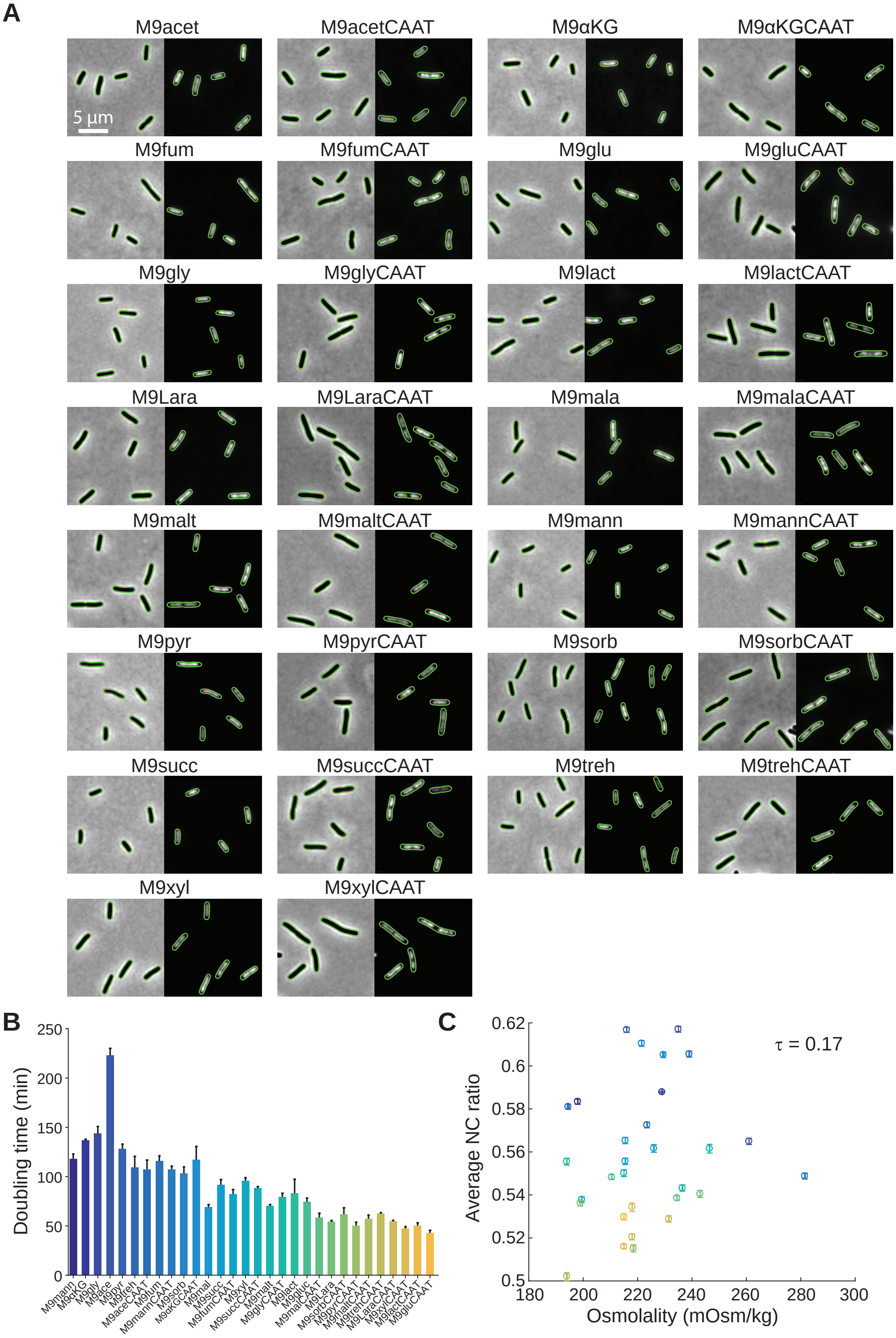
A
M9acet
M9acetCAAT
M9αKG
M9αKGCAAT
5 μm
M9fum
M9fumCAAT
M9glu
M9gluCAAT
M9gly
M9glyCAAT
M9lact
M9lactCAAT
M9Lara
M9LaraCAAT
M9mala
M9malaCAAT
M9malt
M9maltCAAT
M9mann
M9mannCAAT
M9pyr
M9pyrCAAT
M9sorb
M9sorbCAAT
M9succ
M9succCAAT
M9treh
M9trehCAAT
M9xyl
M9xylCAAT
B
Doubling time (min)
250
200
150
100
50
0
M9mann
M9αKG
M9gly
M9ace
M9pyr
M9treh
M9aceCAAT
M9fum
M9mannCAAT
M9sorb
M9αKGCAAT
M9mal
M9succ
M9fumCAAT
M9xyl
M9succCAAT
M9malt
M9glyCAAT
M9lact
M9gluc
M9malCAAT
M9Lara
M9sorbCAAT
M9pyrCAAT
M9maltCAAT
M9trehCAAT
M9LaraCAAT
M9xylCAAT
M9lactCAAT
M9gluCAAT
C
Average NC ratio
0.62
0.6
0.58
0.56
0.54
0.52
0.5
τ = 0.17
180
200
220
240
260
280
300
Osmolality (mOsm/kg)

**Figure S1**

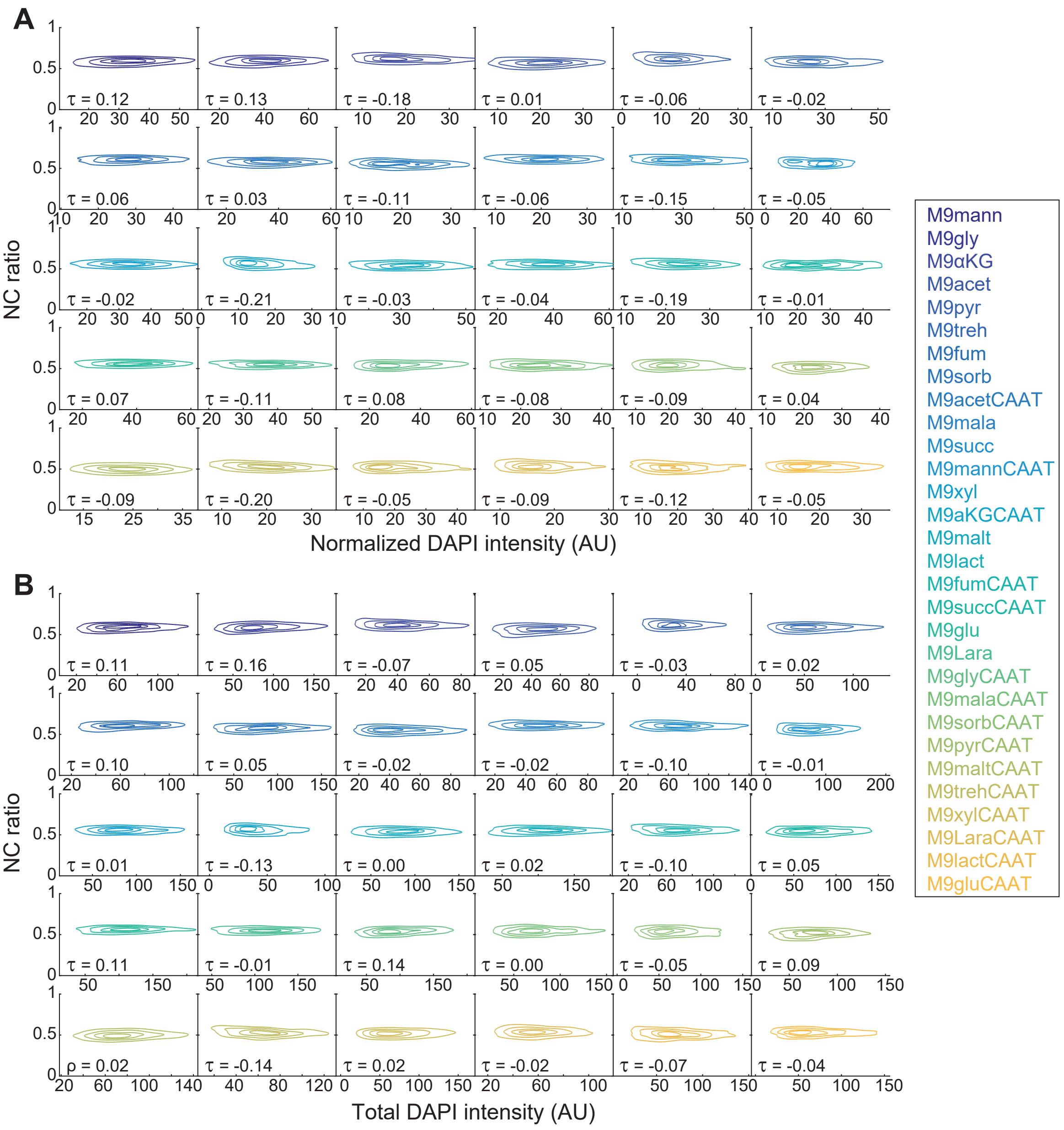
A
NC ratio
Normalized DAPI intensity (AU)
τ = 0.12
τ = 0.13
τ = -0.18
τ = 0.01
τ = -0.06
τ = -0.02
τ = 0.06
τ = 0.03
τ = -0.11
τ = -0.06
τ = -0.15
τ = -0.05
τ = -0.02
τ = -0.21
τ = -0.03
τ = -0.04
τ = -0.19
τ = -0.01
τ = 0.07
τ = -0.11
τ = 0.08
τ = -0.08
τ = -0.09
τ = 0.04
τ = -0.09
τ = -0.20
τ = -0.05
τ = -0.09
τ = -0.12
τ = -0.05
B
NC ratio
Total DAPI intensity (AU)
τ = 0.11
τ = 0.16
τ = -0.07
τ = 0.05
τ = -0.03
τ = 0.02
τ = 0.10
τ = 0.05
τ = -0.02
τ = -0.02
τ = -0.10
τ = -0.01
τ = 0.01
τ = -0.13
τ = 0.00
τ = 0.02
τ = -0.10
τ = 0.05
τ = 0.11
τ = -0.01
τ = 0.14
τ = 0.00
τ = -0.05
τ = 0.09
ρ = 0.02
τ = -0.14
τ = 0.02
τ = -0.02
τ = -0.07
τ = -0.04
M9mann
M9gly
M9αKG
M9acet
M9pyr
M9treh
M9fum
M9sorb
M9acetCAAT
M9mala
M9succ
M9mannCAAT
M9xyl
M9aKGCAAT
M9malt
M9lact
M9fumCAAT
M9succCAAT
M9glu
M9Lara
M9glyCAAT
M9malaCAAT
M9sorbCAAT
M9pyrCAAT
M9maltCAAT
M9trehCAAT
M9xylCAAT
M9LaraCAAT
M9lactCAAT
M9gluCAAT

**Figure S2**

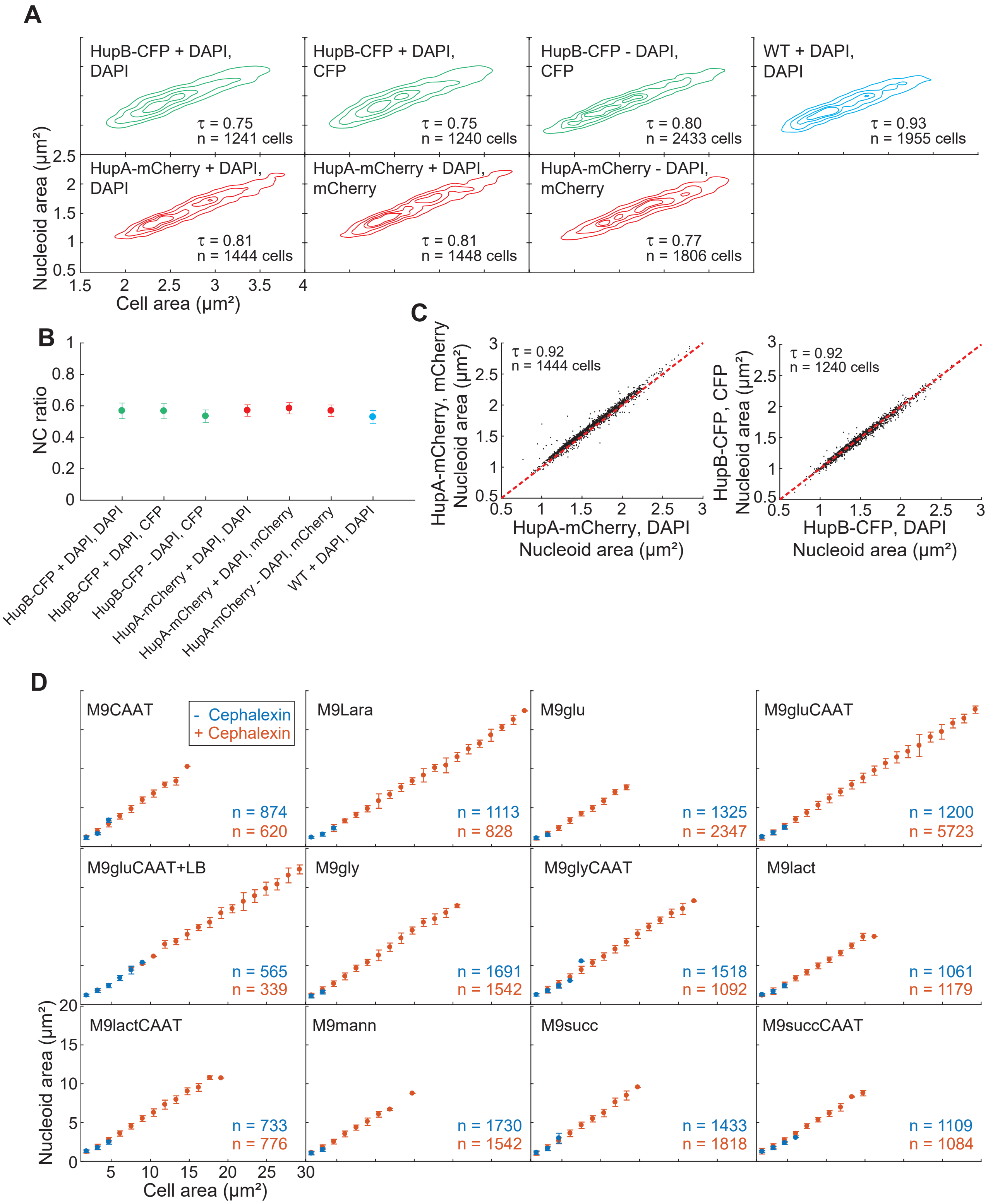
A
HupB-CFP + DAPI, DAPI
τ = 0.75
n = 1241 cells
HupB-CFP + DAPI, CFP
τ = 0.75
n = 1240 cells
HupB-CFP - DAPI, CFP
τ = 0.80
n = 2433 cells
WT + DAPI, DAPI
τ = 0.93
n = 1955 cells
HupA-mCherry + DAPI, DAPI
τ = 0.81
n = 1444 cells
HupA-mCherry + DAPI, mCherry
τ = 0.81
n = 1448 cells
HupA-mCherry - DAPI, mCherry
τ = 0.77
n = 1806 cells
Nucleoid area (μm²)
Cell area (μm²)
B
NC ratio
HupB-CFP + DAPI, DAPI
HupB-CFP + DAPI, CFP
HupB-CFP - DAPI, CFP
HupA-mCherry + DAPI, DAPI
HupA-mCherry + DAPI, mCherry
HupA-mCherry - DAPI, mCherry
WT + DAPI, DAPI
C
τ = 0.92
n = 1444 cells
HupA-mCherry, mCherry Nucleoid area (μm²)
HupA-mCherry, DAPI Nucleoid area (μm²)
τ = 0.92
n = 1240 cells
HupB-CFP, CFP Nucleoid area (μm²)
HupB-CFP, DAPI Nucleoid area (μm²)
D
- Cephalexin
+ Cephalexin
M9CAAT n = 874 n = 620
M9Lara n = 1113 n = 828
M9glu n = 1325 n = 2347
M9gluCAAT n = 1200 n = 5723
M9gluCAAT+LB n = 565 n = 339
M9gly n = 1691 n = 1542
M9glyCAAT n = 1518 n = 1092
M9lact n = 1061 n = 1179
M9lactCAAT n = 733 n = 776
M9mann n = 1730 n = 1542
M9succ n = 1433 n = 1818
M9succCAAT n = 1109 n = 1084
Nucleoid area (μm²)
Cell area (μm²)

**Figure S3**

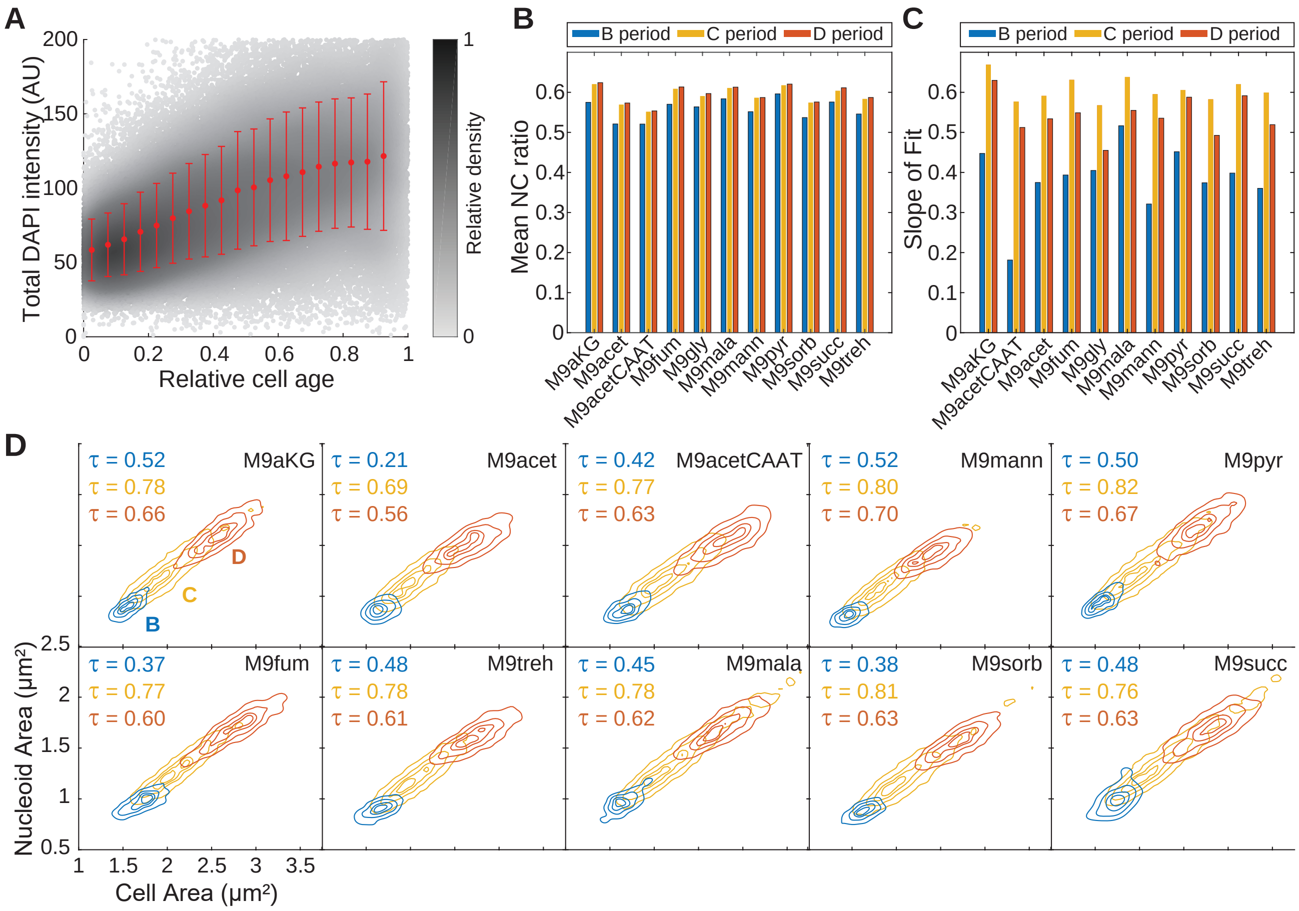
A
Total DAPI intensity (AU)
Relative cell age
Relative density
B
B period
C period
D period
Mean NC ratio
M9aKG
M9acet
M9acetCAAT
M9fum
M9gly
M9mala
M9mann
M9pyr
M9sorb
M9succ
M9treh
C
Slope of Fit
D
τ = 0.52
τ = 0.78
τ = 0.66
M9aKG
τ = 0.21
τ = 0.69
τ = 0.56
M9acet
τ = 0.42
τ = 0.77
τ = 0.63
M9acetCAAT
τ = 0.52
τ = 0.80
τ = 0.70
M9mann
τ = 0.50
τ = 0.82
τ = 0.67
M9pyr
τ = 0.37
τ = 0.77
τ = 0.60
M9fum
τ = 0.48
τ = 0.78
τ = 0.61
M9treh
τ = 0.45
τ = 0.78
τ = 0.62
M9mala
τ = 0.38
τ = 0.81
τ = 0.63
M9sorb
τ = 0.48
τ = 0.76
τ = 0.63
M9succ
Nucleoid Area (μm²)
Cell Area (μm²)

**Figure S4**

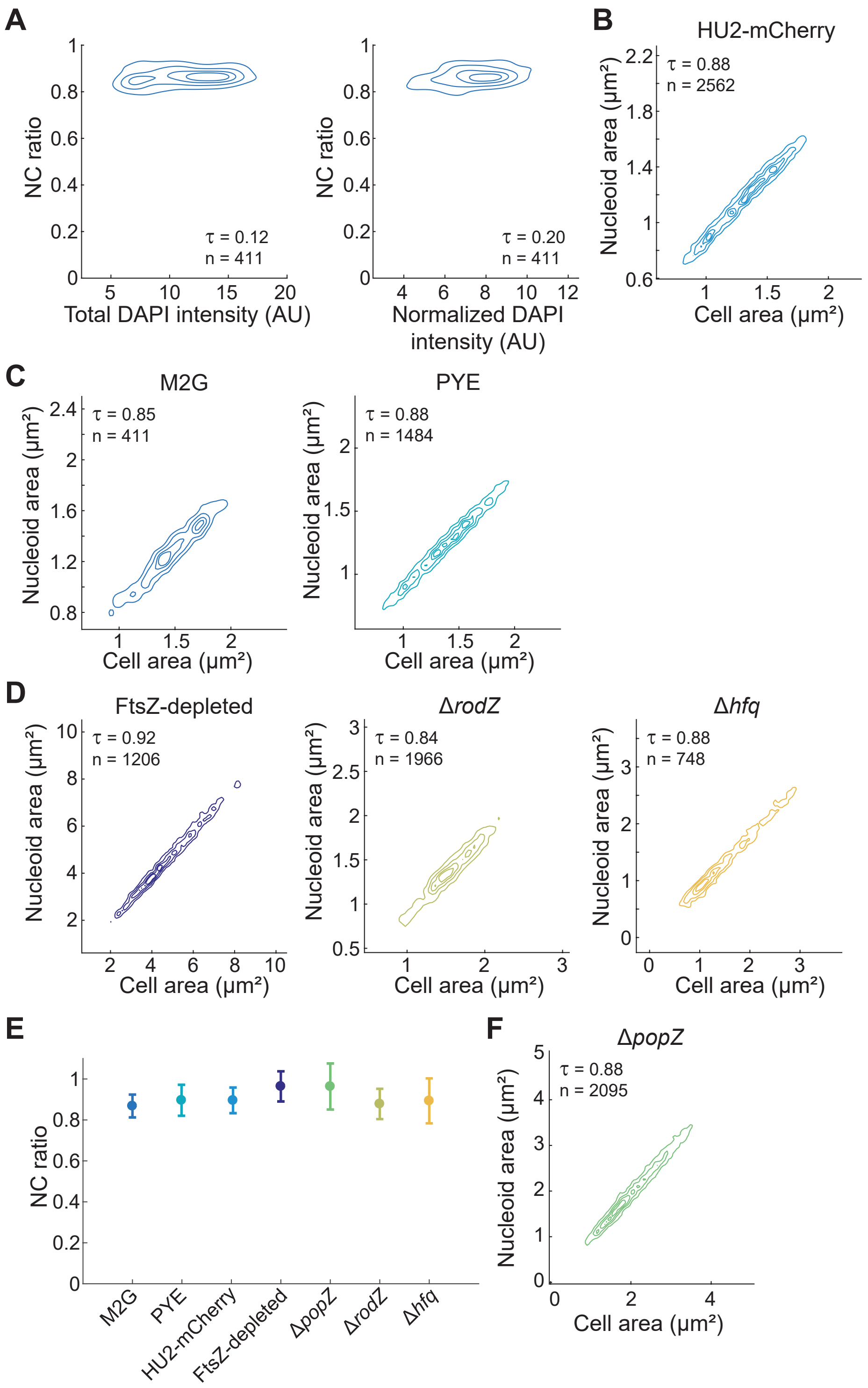
A
NC ratio
τ = 0.12
n = 411
Total DAPI intensity (AU)
τ = 0.20
n = 411
Normalized DAPI intensity (AU)
B
HU2-mCherry
τ = 0.88
n = 2562
Nucleoid area (µm²)
Cell area (µm²)
C
M2G
τ = 0.85
n = 411
PYE
τ = 0.88
n = 1484
D
FtsZ-depleted
τ = 0.92
n = 1206
ΔrodZ
τ = 0.84
n = 1966
Δhfq
τ = 0.88
n = 748
E
M2G
PYE
HU2-mCherry
FtsZ-depleted
ΔpopZ
ΔrodZ
Δhfq
F
ΔpopZ
τ = 0.88
n = 2095

**Figure S5**

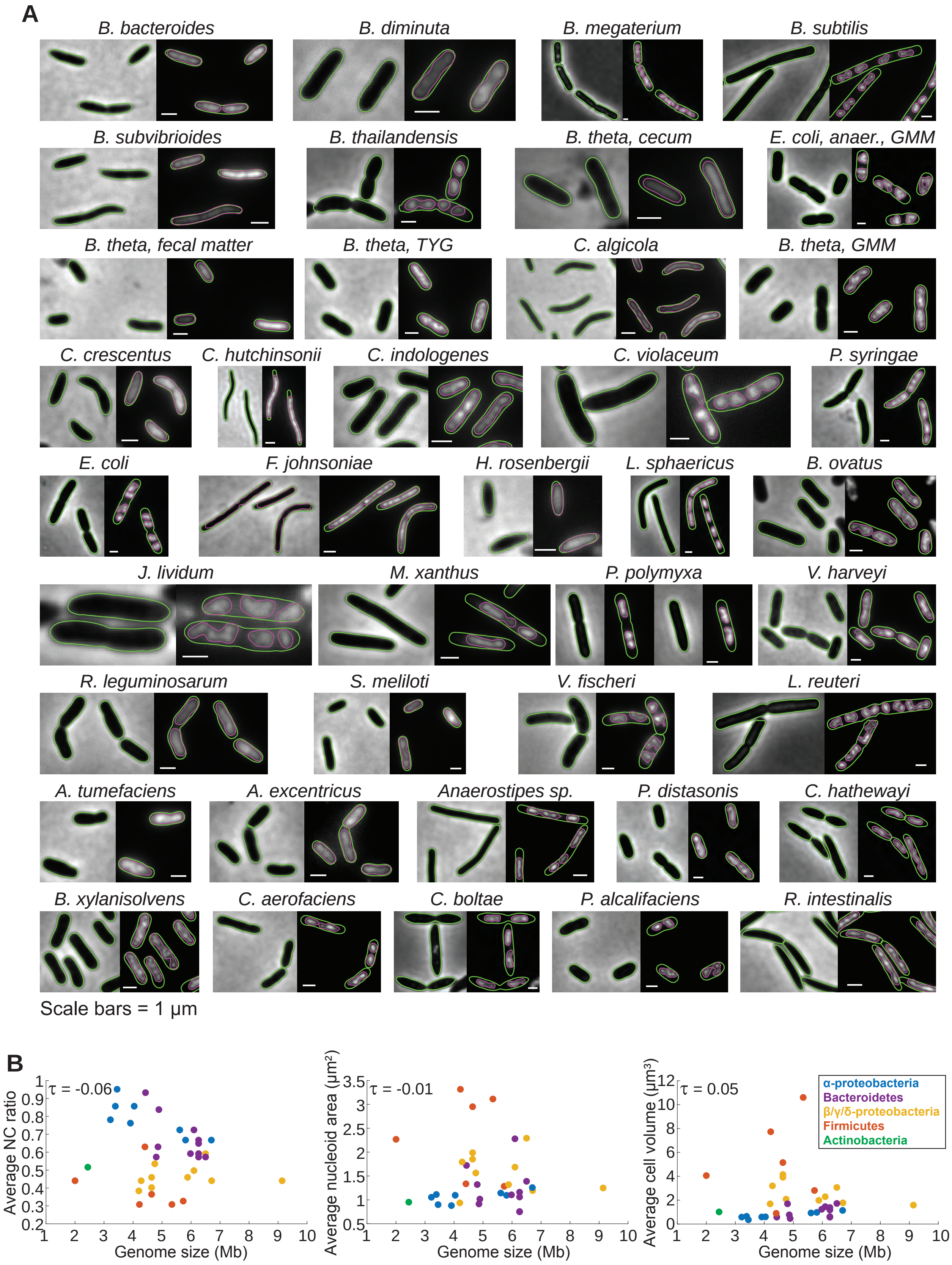
A
B. bacteroides
B. diminuta
B. megaterium
B. subtilis
B. subvibrioides
B. thailandensis
B. theta, cecum
E. coli, anaer., GMM
B. theta, fecal matter
B. theta, TYG
C. algicola
B. theta, GMM
C. crescentus
C. hutchinsonii
C. indologenes
C. violaceum
P. syringae
E. coli
F. johnsoniae
H. rosenbergii
L. sphaericus
B. ovatus
J. lividum
M. xanthus
P. polymyxa
V. harveyi
R. leguminosarum
S. meliloti
V. fischeri
L. reuteri
A. tumefaciens
A. excentricus
Anaerostipes sp.
P. distasonis
C. hathewayi
B. xylanisolvens
C. aerofaciens
C. boltae
P. alcalifaciens
R. intestinalis
Scale bars = 1 μm
B
τ = -0.06
Average NC ratio
Genome size (Mb)
τ = -0.01
Average nucleoid area (μm²)
Genome size (Mb)
τ = 0.05
Average cell volume (μm³)
Genome size (Mb)
α-proteobacteria
Bacteroidetes
β/γ/δ-proteobacteria
Firmicutes
Actinobacteria

**Figure S6**

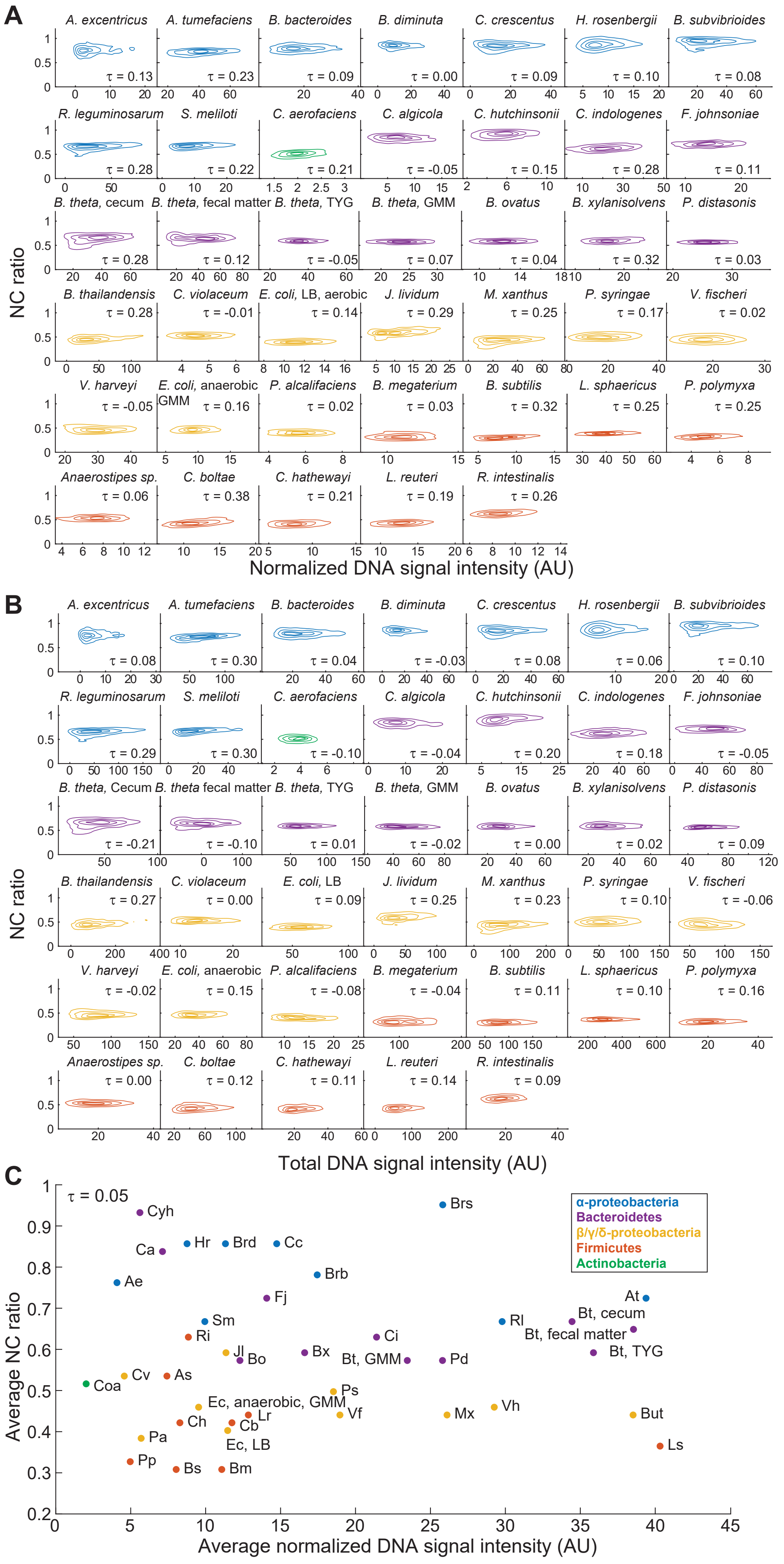
A
B
C
NC ratio
Normalized DNA signal intensity (AU)
Total DNA signal intensity (AU)
Average NC ratio
Average normalized DNA signal intensity (AU)
τ = 0.05
α-proteobacteria
Bacteroidetes
β/γ/δ-proteobacteria
Firmicutes
Actinobacteria

**Figure S7**

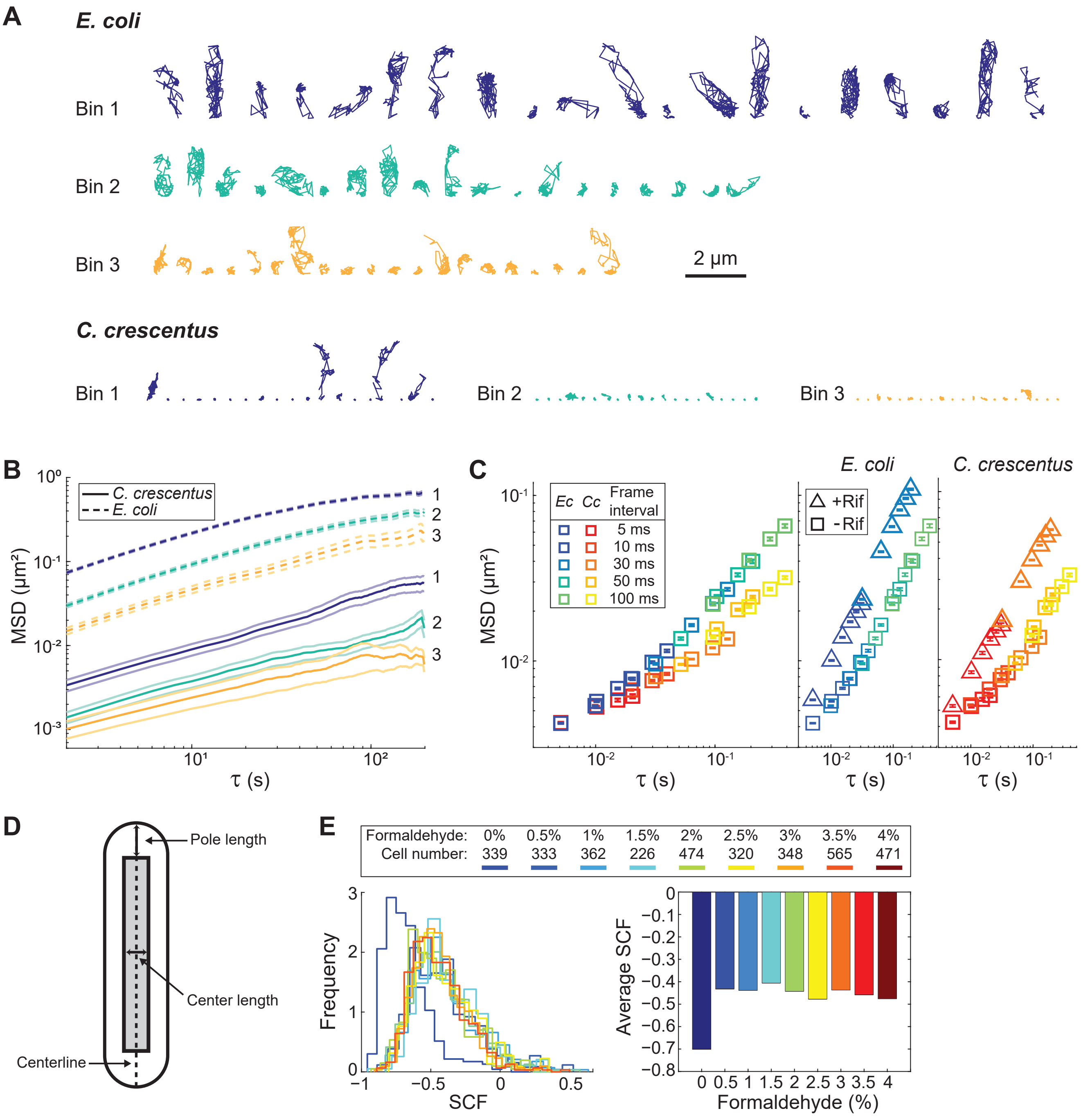
A
E. coli
Bin 1
Bin 2
Bin 3
2 μm
C. crescentus
Bin 1
Bin 2
Bin 3
B
C. crescentus
E. coli
MSD (μm²)
τ (s)
C
E. coli
C. crescentus
Ec Cc Frame interval
5 ms
10 ms
30 ms
50 ms
100 ms
+Rif
-Rif
MSD (μm²)
τ (s)
D
Pole length
Center length
Centerline
E
Formaldehyde: 0% 0.5% 1% 1.5% 2% 2.5% 3% 3.5% 4%
Cell number: 339 333 362 226 474 320 348 565 471
Frequency
SCF
Average SCF
Formaldehyde (%)

**Figure S8**
